# CUT&Tag Identifies Repetitive Genomic Loci that are Excluded from ChIP Assays

**DOI:** 10.1101/2025.02.03.636299

**Authors:** Brandon J Park, Shan Hua, Karli D Casler, Eric Cefaloni, Michael C Ayers, Rahiim F Lake, Kristin E Murphy, Paula M Vertino, Mitchell R. O’Connell, Patrick J Murphy

**Author notes:** Co-corresponding Authorship.

## Abstract

Determining the genomic localization of chromatin features is an essential aspect of investigating gene expression control, and ChIP-Seq has long been the gold standard technique for interrogating chromatin landscapes. Recently, the development of alternative methods, such as CUT&Tag, have provided researchers with alternative strategies that eliminate the need for chromatin purification, and allow for *in situ* investigation of histone modifications and chromatin bound factors. Mindful of technical differences, we set out to investigate whether distinct chromatin modifications were equally compatible with these different chromatin interrogation techniques. We found that ChIP-Seq and CUT&Tag performed similarly for modifications known to reside at gene regulatory regions, such as promoters and enhancers, but major differences were observed when we assessed enrichment over heterochromatin-associated loci. Unlike ChIP-Seq, CUT&Tag detects robust levels of H3K9me3 at a substantial number of repetitive elements, with especially high sensitivity over evolutionarily young retrotransposons. IAPEz-int elements for example, exhibited underrepresentation in mouse ChIP-Seq datasets but strong enrichment using CUT&Tag. Additionally, we identified several euchromatin-associated proteins that co-purify with repetitive loci and are similarly depleted when applying ChIP-based methods. This study reveals that our current knowledge of chromatin states across the heterochromatin portions of the mammalian genome is extensively incomplete, largely due to limitations of ChIP-Seq. We also demonstrate that newer *in situ* chromatin fragmentation-based techniques, such as CUT&Tag and CUT&RUN, are more suitable for studying chromatin modifications over repetitive elements and retrotransposons.

**Highlights:** *In situ* fragmentation overcomes biases produced by ChIP-Seq.

Heterochromatic regions of the genome are lost to the insoluble pellet during ChIP-Seq.

CUT&Tag allows for mapping chromatin features at young repetitive elements.

Euchromatin-associated regulatory factors co-purify with insoluble heterochromatin.

## Introduction

Epigenetic marks and chromatin modifications influence chromatin packaging and regulate gene expression^1,2^. Many of these features are known to play crucial roles in organismal development, and mis-regulation has been associated with a variety of diseases^3–5^. CUT&Tag is a relatively new genomics technique that utilizes a Tn5 transposase to map the genomic location of chromatin modifications^6^. Tn5 allows users to specifically cleave DNA at target genomic locations that are marked by a certain chromatin feature, without the need for crosslinking or sonication^7^. Prior studies have demonstrated that CUT&Tag offers increased specificity, increased signal to noise ratios, requires fewer cells as input, and can be more cost effective than ChIP-Seq^6^, making it an attractive alternative in many situations. While both CUT&Tag and ChIP-Seq are capable of mapping most epigenetic marks, prior studies have uncovered inherent biases caused by the application of ChIP-Seq, potentially limiting investigation of certain chromatin features^8,9^. For example, input material for ChIP-Seq has been found to be biased for open and accessible regions of the genome, and against condensed loci, potentially due to differences in DNA sensitivity to sonication or cross-linking^10,11^. Whether CUT&Tag or CUT&RUN can overcome such biases remains undetermined.

Many heterochromatic regions of the genome contain repetitive elements or retrotransposons, which remain transcriptionally silent in most tissues to prevent spreading of mobile DNA elements throughout the genome, which can cause mutations and DNA damage^12,13^. With recent advances in technology and release of the T2T-CHM13 human genome assembly^14,15^, a renewed emphasis has been placed on the investigation of non-coding DNA sequences, including retrotransposons. Various prior studies have demonstrated that certain retrotransposons play important roles in diverse biological processes, including development, immune response, and neurological function^16–18^. Additionally, aberrant expression of repetitive elements has recently been linked with disease states, including cancer^19^. Thus, establishing a deeper understanding of chromatin states at repetitive elements and retrotransposons is central for advancing biological research across a wide range of fields. Accurately interrogating chromatin states over heterochromatic is essential to facilitate forthcoming research into repetitive element function.

Chromatin features, including post-translational modifications to histones and histone variants, are known to be involved in regulating chromatin packaging and gene expression patterns in countless biological systems^1,2^. Certain modifications and variants have been associated with condensed chromatin and transcriptional repression, while others have been associated with accessible regions of the genome that are actively expressed. H3K9me3 (Histone H3 Lysine 9 trimethylation), for example, is one of the most well studied marks known to reside at constitutive heterochromatin (regions of silent highly compacted DNA), while H3K27me3 (Histone H3 Lysine 27 trimethylation) is primarily found at facultative heterochromatin (regions selectively silenced in specific cell types or developmental stages)^20,21^. The majority of repetitive genomic regions are marked by these repressive modifications in differentiated somatic cells, whereas activating histone modifications can occur when repetitive elements become expressed^16,19^. Activating chromatin features, including H3K27ac (Histone H3 Lysine 27 acetylation) and the histone H2A variant H2A.Z, are typically found over actively expressed euchromatic regions of the genome, such as promoters and enhancers^22^. While some features have been observed both at euchromatin and heterochromatin loci, whether they have roles in both activation and silencing remains largely unknown, particularly at repetitive loci^2,23,24^.

Cognizant of the established limitations of ChIP-Seq^8–11^, we wondered whether newer chromatin profiling methods, such as CUT&Tag, might be more effective for investigating heterochromatic loci and repetitive elements. To investigate this possibility, we began by analyzing equivalent ChIP-Seq and CUT&Tag datasets, measuring the enrichment and genomic localization patterns of four separate chromatin modifications, H2A.Z, H3K27ac, H3K27me3, and H3K9me3. We found similar enrichment profiles were present when comparing analogous ChIP-Seq and CUT&Tag datasets measuring H2A.Z, H3K27ac, H3K27me3, but this was not the case for H3K9me3. Across several distinct mouse and human cell types, measurements of H3K9me3 enrichment were more robust in datasets generated by CUT&Tag than those generated by ChIP-Seq, which facilitated in-depth analysis of repetitive element chromatin states. These initial studies led us to investigate sources of biases in ChIP-based strategies, and to assess whether *in situ* chromatin fragmentation methods could overcome these shortcomings. Our results reveal that the current understanding of chromatin regulation is severely limited due to deficiencies in ChIP-based methods and provide a straightforward route for improved future investigation.

## Results

### ChIP-Seq is Biased in Favor of Gene Promoters and Against Intergenic Regions

To identify genomic loci which might be preferentially enriched in ChIP-Seq datasets, as explored by others previously^10^, we randomly sub-sampled the genome (100,000 1Kb randomly selected regions) and partitioned regions into quartiles based on normalized enrichment scores (RPKM) from publicly available ChIP-Seq data, generated from input samples (soluble sonicated chromatin extracted prior to immunoprecipitation) (GEO Accession GSE181069)^25^ purified from mouse embryonic fibroblasts (MEFs). Using standard peak-calling strategies for identifying enriched regions (see methods), we partitioned the top 20,000 genomic regions possessing the highest ChIP-Seq input enrichment scores, termed ‘Top Input’, and assessed overall genomic context. Loci with high relative input scores (putative ChIP-Seq false positives) (Quartile 4 and Top Input regions) were located in closer proximity to gene transcription start sites (TSSs) than regions with lower enrichment scores (Quartile 1) (**Fig 1A**), and these regions included a relatively large number of gene promoters (**Fig 1B**). They also possessed differing levels of CpG density (**Supp Fig 1A**) and had highly accessible chromatin, as measured by ATAC-Seq^26^ (GEO Accession GSE145705) (**Fig 1C**). In agreement with these observations, enrichment values for ChIP-Seq input were highly correlated with chromatin accessibility measurements at gene promoters (R=0.76) (**Fig 1D, 1E, & Supp Fig 1B**). Taken together, these results align with prior studies which report biases from ChIP, with potential artifacts caused by a preferential selection of euchromatin at the expense of heterochromatic loci^10^.

**Figure 1.**
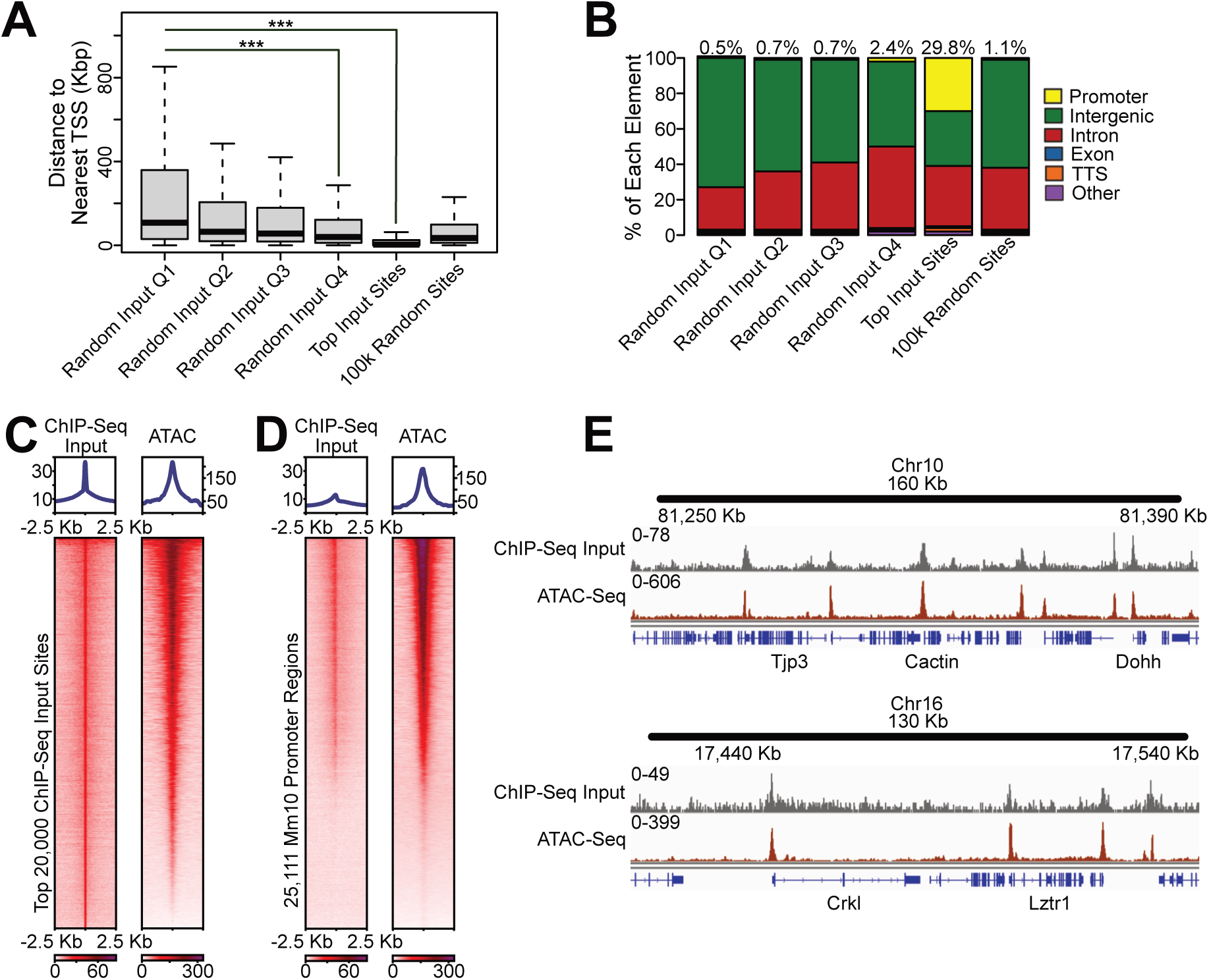
ChIP-Seq Input Samples are Enriched for Promoters and Open Chromatin. (A) Distance to nearest gene transcription start site (TSS) for ChIP-Seq input (100k random 1Kb regions divided into quartiles based on input enrichment levels), top 20k ChIP-Seq input sites, or 100k random 1Kb sites. (B) Genomic annotation for ChIP-Seq input (100k random 1Kb regions divided into quartiles based on input enrichment levels), top 20k ChIP-Seq input sites, or 100k random 1Kb sites. Percentages indicate % of promoter regions. (C) Heatmaps and profile plots of enrichment scores (RPKM) for ChIP-Seq input and ATAC-Seq datasets over top 20k input sites. (D) Heatmaps of enrichment scores (RPKM) for ChIP-Seq input and ATAC-Seq datasets over all annotated mouse promoters. (E) Genome browser enrichment profiles of ChIP-Seq input and ATAC-Seq showing overlap between the methods.

### ChIP-Seq and CUT&Tag Display Similar Enrichment Patterns for Activating Chromatin Marks

Because *in situ* chromatin profiling methods, such as CUT&Tag, are methodologically distinct from immunoprecipitation-based techniques, we next wondered whether enrichment profiles generated by CUT&Tag differed from profiles generated by ChIP-Seq. Despite the potential biases of ChIP-Seq, effective enrichment scores could be attained by comparing DNA isolated from immunoprecipitated material with DNA isolated from input samples (scored as -log_10_ p-values from a Poisson distribution). Using this approach, we compared separate enrichment profiles, and began by investigating the activating chromatin modifications H2A.Z^27^ (GEO Accession GSE51579) and H3K27ac^28^ (GEO Accession GSE72239). CUT&Tag replicates were consistent for chromatin features (**Supp Fig 2A**), and similar enrichment patterns were observed over gene promoters and highly enriched regions (peaks) regardless of technique (**Fig. 2A & 2B**). Likewise, the most highly enriched regions of the genome for both H2A.Z and H3K27ac tended to occur in close proximity to gene transcription start sites (**Fig. 2C),** and these features were found to be preferentially located within gene regulatory regions such as promoters (**Fig. 2D)**. Finally, to directly compare enrichment profiles for ChIP-Seq with profiles from CUT&Tag (and overcome potential differences in data processing), we rank normalized and assessed the degree of correlation between datasets. As anticipated, rank scores over highly enriched loci and gene promoters were found to be well correlated (**Fig. 2E & Supp Fig 2B**). These results demonstrate that ChIP-Seq and CUT&Tag perform similarly for the activating chromatin marks H2A.Z and H3K27ac, which are known to preferentially reside over gene promoters and transcriptionally active gene regulatory regions ^2,22,23^.

**Figure 2.**
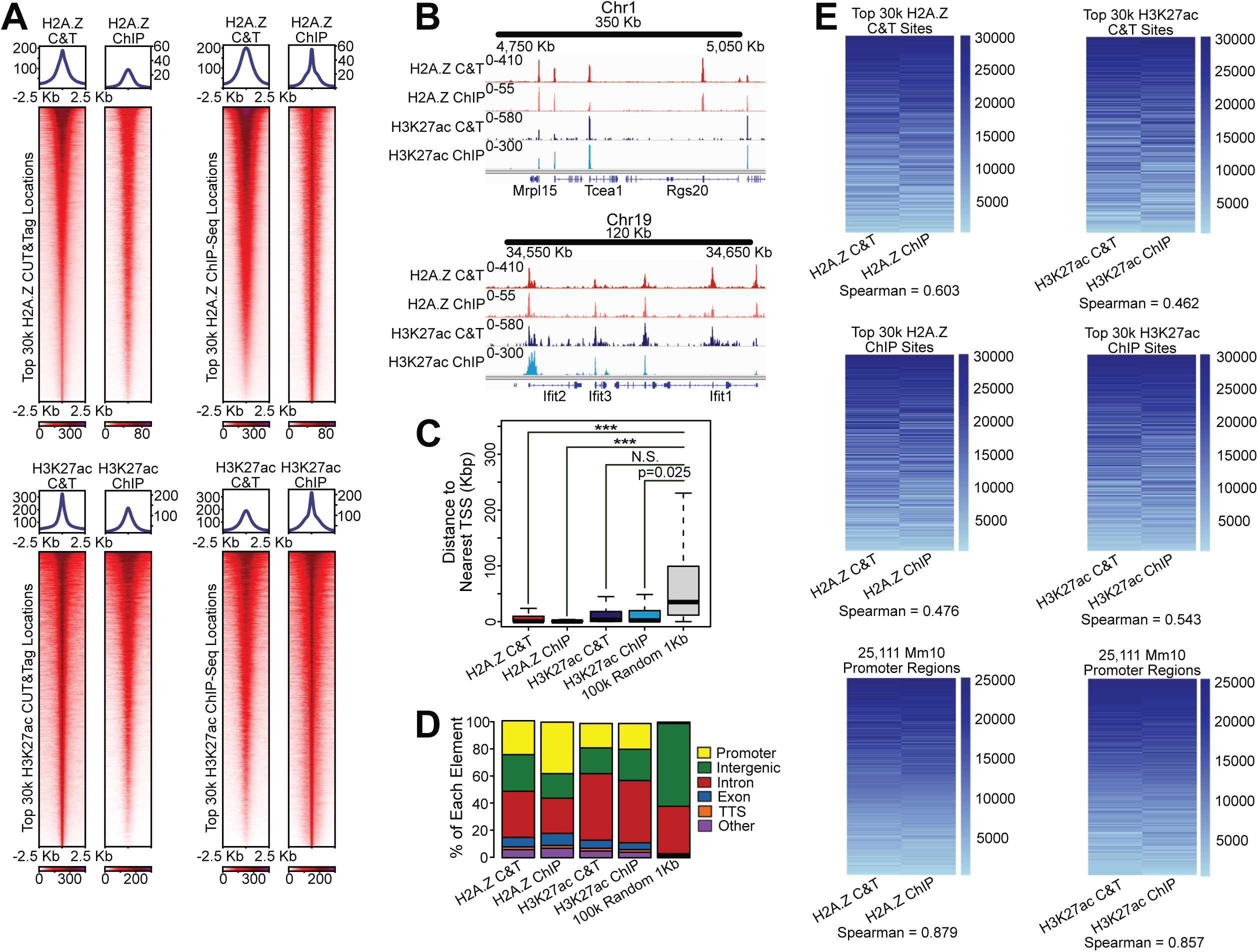
ChIP-Seq and CUT&Tag Produce Similar H2A.Z and H3K27ac Chromatin Landscapes. (A) Heatmaps and profile plots of enrichment scores from H2A.Z and H3K27ac CUT&Tag and ChIP-Seq datasets. H2A.Z datasets were plotted over the top 30k H2A.Z CUT&Tag and ChIP-Seq sites, and H3K27ac datasets were plotted over the top 30k H3K27ac CUT&Tag and ChIP-Seq sites. (B) Genome browser enrichment profiles of H2A.Z and H3K27ac CUT&Tag and ChIP-Seq datasets showing overlap between the methods for both chromatin marks. (C) Distance to nearest gene transcription start site for 30k most highly enriched H2A.Z CUT&Tag sites, 30k most highly enriched H2A.Z ChIP-Seq sites, 30k most highly enriched H3K27ac CUT&Tag sites, 30k most highly enriched H3K27ac ChIP-Seq sites, or 100k random 1Kb sites. (D) Genomic annotation for 30k most highly enriched H2A.Z CUT&Tag sites, 30k most highly enriched H2A.Z ChIP-Seq sites, 30k most highly enriched H3K27ac CUT&Tag sites, 30k most highly enriched H3K27ac ChIP-Seq sites, or 100k random 1Kb sites. (E) Heatmaps of rank normalized enrichment scores for H2A.Z and H3K27ac CUT&Tag and ChIP-Seq datasets. H2A.Z datasets were plotted over the top 30k H2A.Z CUT&Tag sites, the top 30k H2A.Z ChIP-Seq sites, and 100k random 1Kb regions. H3K27ac datasets were plotted over the top 30k H3K27ac CUT&Tag sites, the top 30k H3K27ac ChIP-Seq sites, and 100k random 1Kb regions.

### ChIP-Seq and CUT&Tag Display Dissimilar Enrichment Patterns for the Repressive Chromatin Mark H3K9me3

Considering the preferential enrichment we observed over gene promoters for ChIP-Seq input samples (**Fig 1**) and our putative ability to overcome these biases (by comparing immunoprecipitated DNA and input DNA) (**Fig 2**), we next reasoned that chromatin features located outside of gene promoters, within intergenic regions, may be inadvertently excluded from ChIP-Seq studies. To investigate this possibility, we focused our analysis on chromatin modifications that are located primarily within facultative and constitutive heterochromatin, H3K27me3^28^ (GEO Accession GSE72239) and H3K9me3 (GEO Accession GSE181069), respectively. As with our prior comparisons, individual CUT&Tag replicates were highly similar for H3K27me3 and H3K9me3 (**Supp Fig 3A**), and highly enriched H3K27me3 sites (top 10,000) were identified by both techniques (**Fig 3A-3B & Supp Fig 3B**). These highly enriched H3K27me3 sites tended to occur in close proximity to gene transcription start sites, and similar types of genomic locations were enriched regardless of technique (**Fig 3C & 3D**). However, regions highly enriched for H3K9me3 (top 30,000) were largely distinct when comparing between ChIP-Seq and CUT&Tag (**Fig 3A & 3B).** Although both techniques identified a similar subset of genomic locations, regions identified as enriched using ChIP-Seq were located in closer proximity to gene transcription start sites than analogous regions identified by CUT&Tag (**Fig 3C & 3D**). To assess correlation between data from separate methods, we again performed rank normalization and found H3K27me3 scores to be moderately correlated (**Fig 3E & Supp Fig 3C**). In contrast, H3K9me3 scores displayed particularly low correlations at the most highly enriched CUT&Tag and ChIP-Seq sites (R=0.128 and 0.152 respectively). Similar to the activating marks, the highest correlation values for H3K27me3 were observed over gene promoter regions, while there was an inverse correlation over promoters for H3K9me3 (R=-0.241). These results indicate that CUT&Tag and ChIP-Seq perform similarly well for the repressive H3K27me3 modification, but the two techniques produce very different enrichment profiles for the repressive H3K9me3 modification, which is typically found at silent heterochromatic loci.

**Figure 3.**
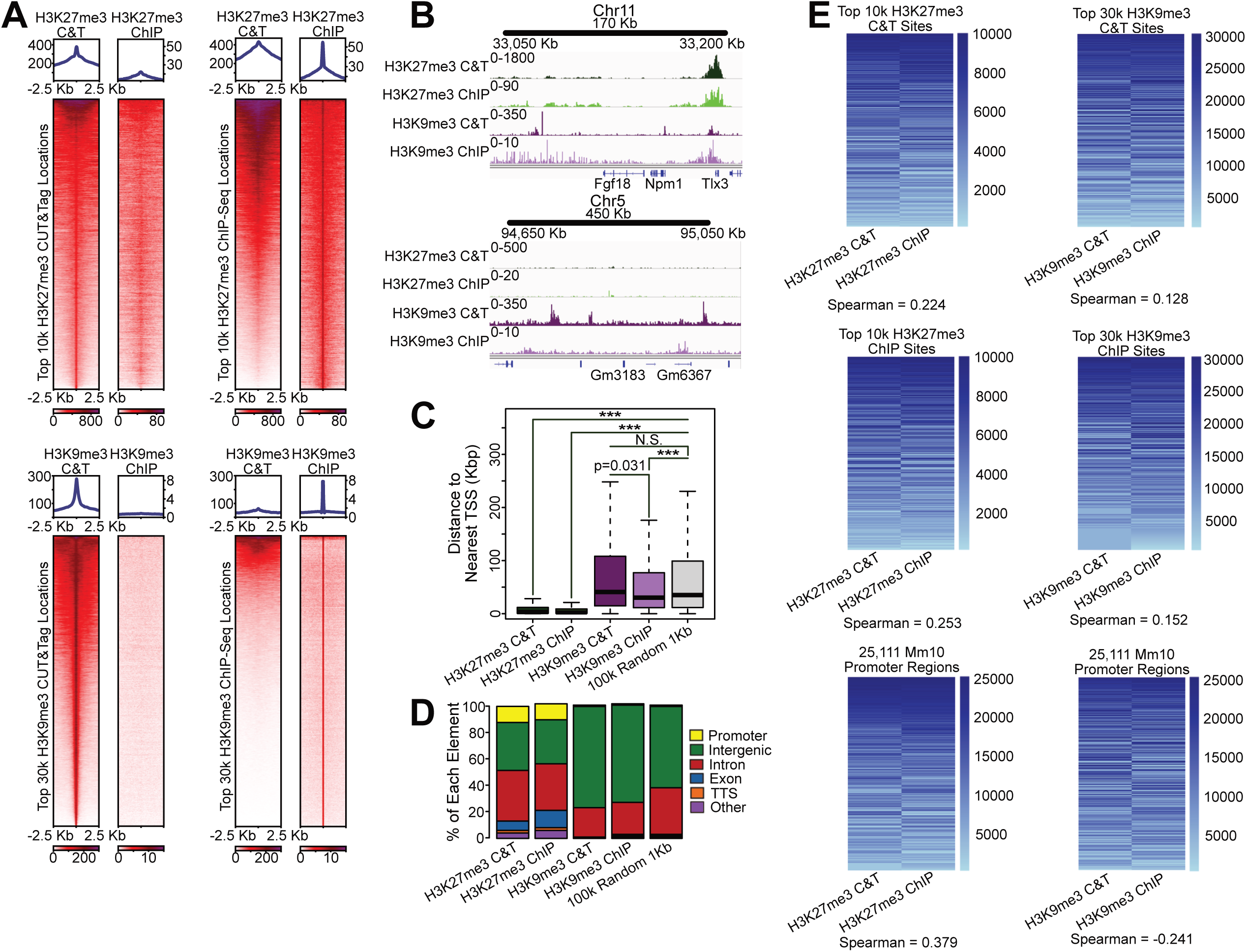
ChIP-Seq and CUT&Tag Produce Similar Chromatin Landscapes for H3K27me3, but Not H3K9me3. (A) Heatmaps and profile plots of enrichment scores from H3K27me3 and H3K9me3 CUT&Tag and ChIP-Seq datasets. H3K27me3 datasets were plotted over the top 10k H3K27me3CUT&Tag and ChIP-Seq sites, and H3K9me3 datasets were plotted over the top 30k H3K9me3 CUT&Tag and ChIP-Seq sites. (B) Genome browser enrichment profiles of H3K27me3 and H3K9me3 CUT&Tag and ChIP-Seq datasets showing overlap between the methods for H3K27me3, but not H3K9me3. (C) Distance to nearest gene transcription start site for 30k most highly enriched H3K27me3 CUT&Tag sites, 30k most highly enriched H3K27me3ChIP-Seq sites, 30k most highly enriched H3K9me3 CUT&Tag sites, 30k most highly enriched H3K9me3 ChIP-Seq sites, or 100k random 1Kb sites. (D) Genomic annotation for 30k most highly enriched H3K27me3 CUT&Tag sites, 30k most highly enriched H3K27me3ChIP-Seq sites, 30k most highly enriched H3K9me3 CUT&Tag sites, 30k most highly enriched H3K9me3 ChIP-Seq sites, or 100k random 1Kb sites. (E) Heatmaps of rank normalized enrichment scores for H3K27me3 and H3K9me3 CUT&Tag and ChIP-Seq datasets. H3K27me3 datasets were plotted over the top 10k H3K27me3CUT&Tag sites, the top 10k H3K27me3 ChIP-Seq sites, and 100k random 1Kb regions. H3K9me3 datasets were plotted over the top 30k H3K9me3 CUT&Tag sites, the top 30k H3K9me3 ChIP-Seq sites, and 100k random 1Kb regions.

### Crosslinking and Sonication Create an Over-Representation of Euchromatin and an Under-Representation of Intergenic Heterochromatin

Potential biases in ChIP-Seq might arise through increased sensitivity to sonication at euchromatic loci, by a resistance to sonication at heterochromatic loci, or a combination of both factors. To investigate these possibilities, we performed a mock ChIP-Seq experiment on wild-type primary MEFs, sonicating chromatin to varying degrees and isolating DNA from the soluble fraction, which is commonly used for ChIP experiments, and the insoluble fraction, which is typically discarded. We then compared the DNA purified from each mock ChIP-Seq sample. DNA fragments from insoluble pellet samples (from cross-linked minimally sonicated chromatin) exhibited a much larger size (higher molecular weight) than DNA from soluble supernatant fractions (**Fig 4A**). To establish whether distinct portions of the genome reside within these separate fractions (potentially underlying biases in ChIP-Seq), we performed Illumina sequencing on isolated DNA, including the minimally sonicated soluble chromatin (Cross-linked Sonicated Supernatant 1 – S1), thoroughly sonicated soluble chromatin (Cross-linked Sonicated Supernatant 3 – S3), and insoluble chromatin (Cross-linked Sonicated Pellet 1 – P1). Similar to our observations from ChIP-Seq input sample measurements, DNA isolated from minimally (S1) and thoroughly (S3) sonicated soluble chromatin was found to be enriched over euchromatic gene promoters, while DNA from the insoluble pellet (P1) was more enriched over intergenic regions (**Fig 4B**). Additionally, highly enriched minimally sonicated supernatant DNA (S1) tended to localize in close proximity to gene TSSs (**Fig 4C**), and enriched regions from both supernatant samples (S1 and S3) had high CpG densities (**Fig 4D**). Higher levels of enrichment were also detected over gene promoters when comparing between the supernatant and insoluble pellet samples (**Fig 4E**), as well as regions previously identified in Figure 1 as enriched in ChIP-Seq input datasets (**Fig 4F**).

**Figure 4.**
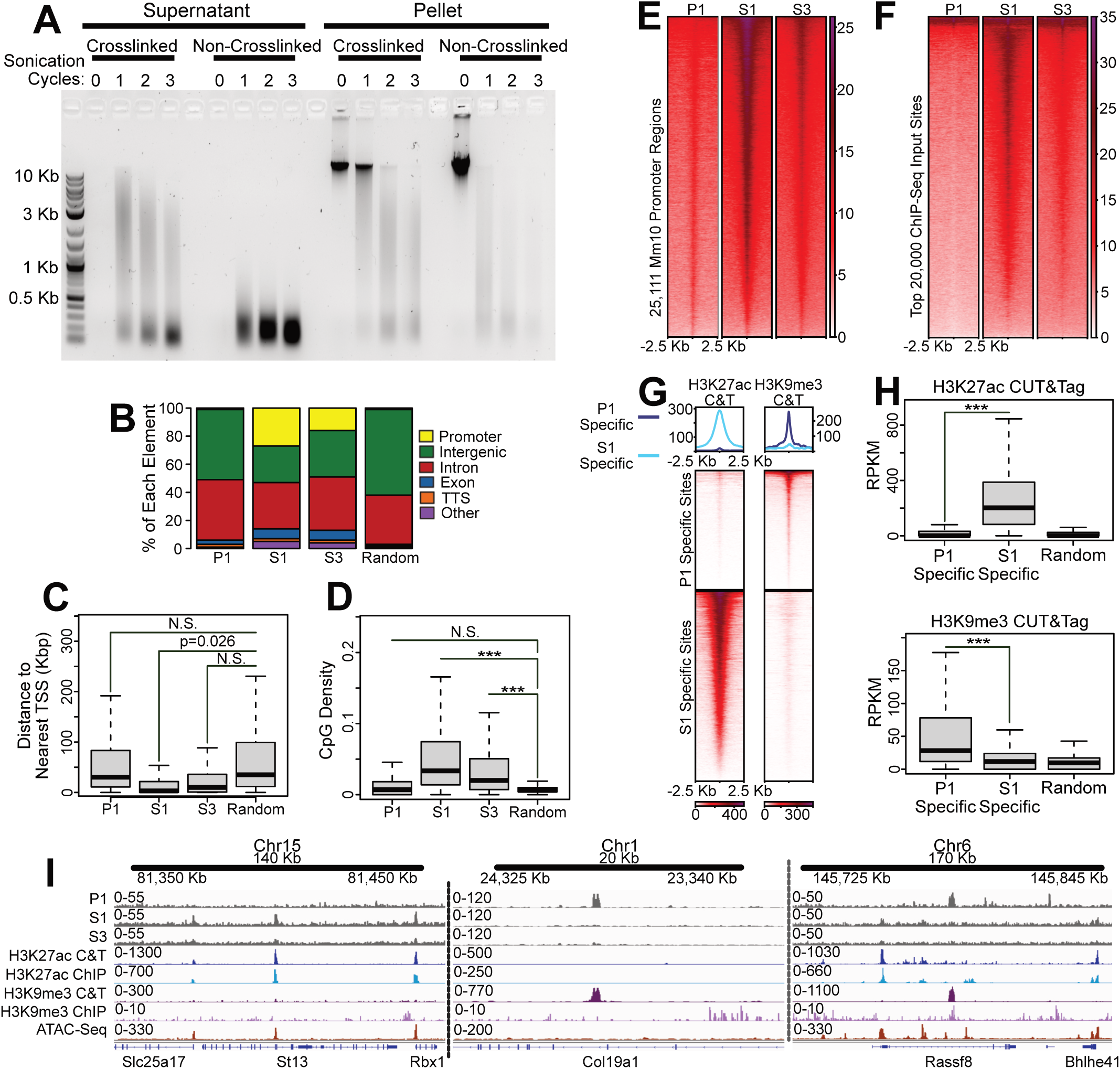
Crosslinked and Sonicated Soluble Chromatin is Enriched for Promoters, and Insoluble Chromatin is Enriched for Heterochromatic Regions. (A) Agarose gel showing DNA extracted from supernatant or cellular debris pellet after a mock ChIP-Seq input experiment. Samples were crosslinked or left as non-crosslinked controls, and sonicated for 0-3 cycles. (B) Genomic annotation of the top 30k most highly enriched P1, S1, or S3 sites, and 100k random 1Kb regions. (C) Distance to the nearest gene transcription start site of the top 30k most highly enriched P1, S1, or S3 sites, and 100k random 1Kb regions. (D) CpG density of the top 30k most highly enriched P1, S1, or S3 sites, and 100k random 1Kb regions. (E) Heatmaps of P1, S1, and S3 datasets over all the annotated mouse promoters. (F) Heatmaps of P1, S1, and S3 datasets over the top 20k ChIP-Seq input sites identified in Figure 1. (G) Heatmaps and profile plots of H3K27ac and H3K9me3 CUT&Tag signal over the most highly enriched P1-specific and S1-specific regions. (H) Enrichment scores of H3K27ac and H3K9me3 CUT&Tag signal over the most highly enriched P1-specific sites, the most highly enriched S1-specific sites, and 100k random 1Kb regions. (I) Genome browser enrichment profiles of P1, S1, and S3 datasets with H3K27ac CUT&Tag, H3K27ac ChIP-Seq, H3K9me3 CUT&Tag, H3K9me3 ChIP-Seq, and ATAC-Seq.

We next classified genomic loci based on whether they were purified from the soluble or insoluble pellet samples (termed S1-specific or P1-specific, respectively) (see methods). In support of our prior observations (**Fig 1**), we found that the activating histone modification H3K27ac was enriched at S1-specific loci, whereas the repressive histone modification H3K9me3 was enriched at P1-specific loci (**Fig 4G – 4I**). The most highly enriched sites from the soluble chromatin samples were also enriched for H3K27ac, while the most highly enriched sites from the insoluble pellet were enriched for H3K9me3 (**Supp Fig 4A**). Notably, enrichment scores from samples generated by micrococcal nuclease (MNase)-based genomic fragmentation^29^ (GEO Accession GSE153939), which is commonly utilized in native ChIP-Seq^7,30^ and CUT&RUN^31^ experiments, were statistically significant but only moderately different in magnitude (median RPKM = 3.113 and 3.468 respectively) comparing between S1 and P1-specific regions, suggesting that MNase-based methods do not suffer from the same biases as standard ChIP-Seq approaches (**Supp Figs 4B & 4C**). Together these results indicate that biases in our mock ChIP-Seq experiment arose due to the combination of open/accessible genomic loci being over-represented in the soluble fraction and inaccessible/heterochromatic loci being over-represented in the insoluble pellet fraction.

### CUT&Tag Identifies H3K9me3 at Young Repetitive Elements that are Undetectable by ChIP-Seq

Our mock ChIP experiments indicated that inaccessible intergenic loci tend to be preferentially excluded from ChIP-Seq assays (**Fig 4**), potentially explaining the dramatic differences we observed when comparing H3K9me3 patterns obtained from ChIP-Seq with results obtained from CUT&Tag (**Fig 3**). We next speculated that specific genomic loci might be particularly sensitive to these biases, rendering them undetectable by ChIP-Seq and only detectible by CUT&Tag. To identify such regions, we performed *k*-means clustering on a combined set of regions identified as enriched in either CUT&Tag or ChIP-Seq. We identified three discrete clusters, including two with higher H3K9me3 enrichment levels from CUT&Tag (Clusters 1 and 2 – C1 & C2) and one with higher H3K9me3 levels from ChIP-Seq (Cluster 3 – C3) (**Fig 5A**). No such differences were observed when we applied an analogous clustering strategy to analyze H3K27ac, H2A.Z, or H3K27me3, reinforcing our earlier results (**Supp Fig 5A**). Prior studies have demonstrated that intergenic repetitive elements and retrotransposons are commonly marked by H3K9me3^11–13,20^. Interestingly, both C1 and C2 clusters in our H3K9me3 comparisons (regions with high enrichment levels in CUT&Tag) possessed a high abundance of LTR family retrotransposons (**Fig 5B**). To gain insight into which specific LTR transposons might be impacted by ChIP biases, we next performed rank scoring of all uniquely named repetitive elements in the mouse genome, and then subtracted ChIP-Seq rank scores from CUT&Tag rank scores, resulting in a single value for each uniquely named repetitive element. Elements receiving a strong negative score possessed high levels of H3K9me3 specifically in ChIP-Seq datasets, whereas regions with a strong positive score possessed high levels of H3K9me3 specifically in CUT&Tag. We also assessed the evolutionary age of each repetitive element types through the use of milliDiv scoring (base mismatches from the consensus repeat sequence in parts per thousand), with lower scores indicating younger elements^16,32^. Strikingly, we found that the majority of young LTR class repetitive elements exhibited very high levels of H3K9me3 specifically in CUT&Tag, including IAPEz-int, RLTR6-int, and RLTR6B elements. Although many LINEs, such as L1Md_F2, possessed higher ChIP-Seq rank scores, they exhibited a lack of H3K9me3 enrichment in both ChIP-Seq and CUT&Tag (**Fig 5C & 5D**). These results indicate that CUT&Tag is capable of identifying H3K9me3 at specific classes of young repetitive elements that have traditionally been underrepresented in ChIP-Seq datasets.

**Figure 5.**
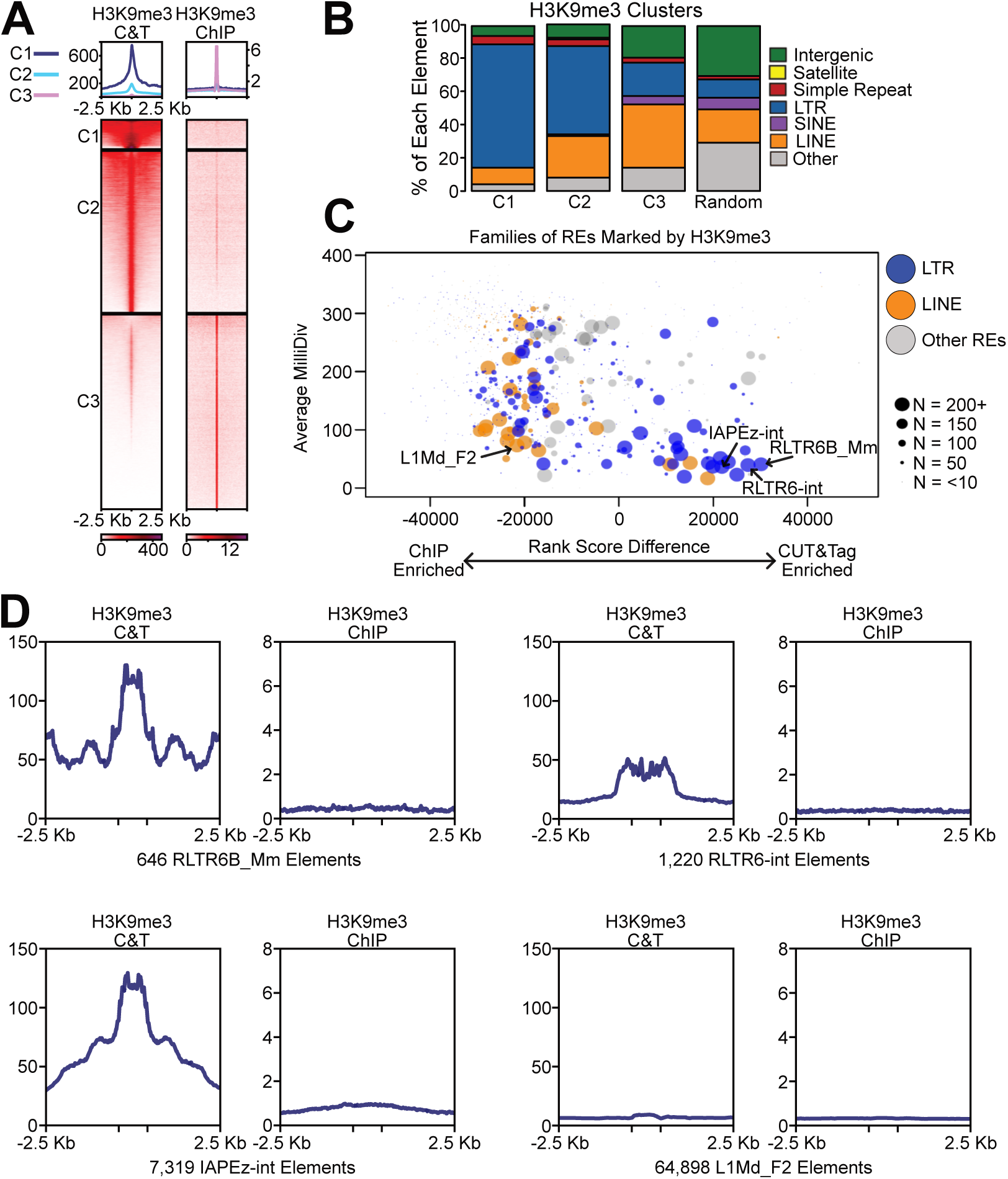
CUT&Tag Identifies H3K9me3 at Evolutionarily Young LTRs. (A) Heatmaps and profile plots of H3K9me3 CUT&Tag and ChIP-Seq datasets over a union file of all the most highly enriched CUT&Tag and ChIP-Seq H3K9me3 sites, sorted by *k*-means clustering (C1-C3). (B) Genomic annotation of the repetitive elements enriched in each cluster (C1-C3). (C) Rank score plot depicting enrichment of various repetitive element families in H3K9me3 CUT&Tag and ChIP-Seq. LTR class elements are labeled in blue, and LINE class elements are labeled in orange. Number of repetitive elements identified in either H3K9me3 dataset are depicted with various sized points based on their abundance in these datasets. (D) Profile plots of H3K9me3 CUT&Tag and ChIP-Seq datasets over all RLTR6B, RLTR6-int, IAPEz-int, and L1Md_F2 elements.

### CUT&Tag Identifies H3K9me3 and H2A.Z at Young Repetitive Elements in Various Mouse and Human Cell Lines

To establish additional support for our findings (**Fig 5**), we repeated our prior analyses using an additional MEF H3K9me3 ChIP-Seq dataset^33^ (GEO Accession GSE53939), as well as CUT&Tag^34^ (GEO Accession GSE213350) and ChIP-Seq (ENCODE ENCSR000APZ) data generated from human H1 stem cells. In all cases, we found young LTR class transposons possessed higher levels of H3K9me3 in CUT&Tag datasets than in ChIP-Seq (**Fig 6A & Supp Figs 6A-6D**). We next compared H3K9me3 CUT&Tag data from MEFs with H3K9me3 CUT&Tag data from mouse embryonic stem cells (mESCs) and H3K9me3 CUT&RUN data from MEFs. Here again, specific classes of evolutionarily young repetitive elements, particularly LTRs, were more highly enriched for H3K9me3 than many of the evolutionarily older elements (**Fig 6B & 6C**). As in our prior results, enrichment for H3K9me3 over IAPEz-int and RLTR6-int elements was particularly highly in the MEF and mESC CUT&Tag datasets, as well as the MEF CUT&RUN dataset. Taken together, these results indicate that *in situ* fragmentation-based methods (such as CUT&Tag or CUT&RUN) can efficiently map many repetitive elements across a variety of cell types, and deficiencies from ChIP-Seq can be effectively overcome with these more recently developed techniques. While discrepancies between ChIP-Seq and CUT&Tag methods were initially identified through measurements of H3K9me3, the possibility remained that additional chromatin features may be present over repetitive elements, such as young LTR transposons, but they have been largely unexplored due to biases of ChIP-Seq. To investigate this possibility, we returned to our prior measurements of H3K27ac, H2A.Z, and H3K27me3. Remarkably, we found that IAPEz-int possessed moderate levels of H2A.Z in CUT&Tag datasets (**Fig 6D & 6E**). Taken together, these results provide compelling evidence that heterochromatic loci and repetitive elements are restricted to the insoluble chromatin fraction during standard ChIP-Seq experiments, that chromatin profiling methods which utilize *in situ* chromatin fragmentation are able to overcome these biases, and that our current knowledge of DNA binding proteins or chromatin modifications localized within heterochromatin regions (such as LTR elements) is decidedly incomplete.

**Figure 6.**
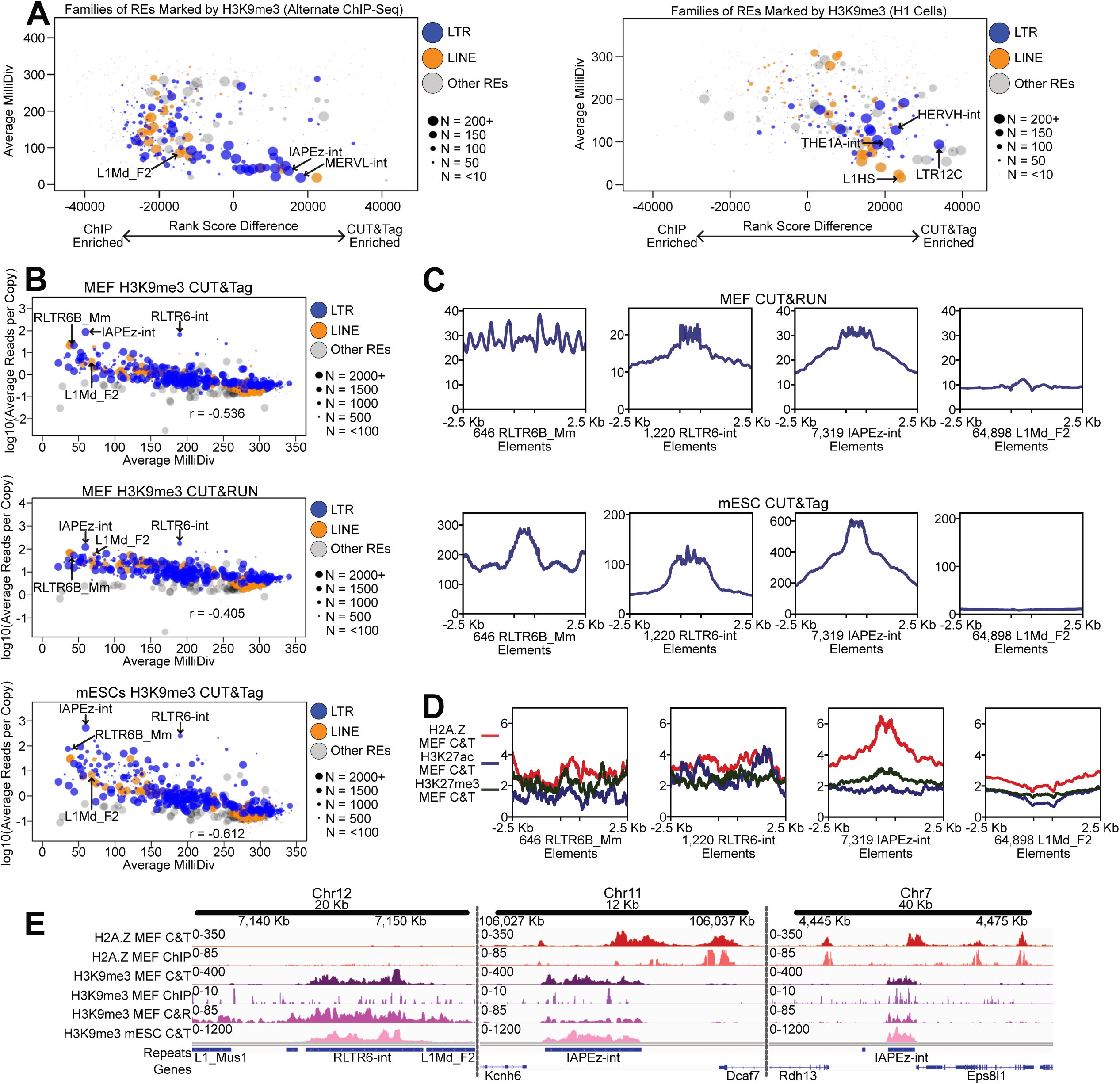
CUT&Tag and CUT&RUN Identify H3K9me3 and H2A.Z at Young Repetitive Elements in Multiple Cell Types. (A) Rank score plots depicting enrichment of various repetitive element families in MEF H3K9me3 CUT&Tag (alternate dataset from Pederson et al.) and ChIP-Seq, as well as H3K9me3 CUT&Tag and ChIP-Seq datasets from human H1 cells. LTR class elements are labeled in blue, and LINE class elements are labeled in orange. Number of repetitive elements identified in either H3K9me3 dataset are depicted with various sized points based on their abundance in these datasets. (B) Scatter plots comparing average milliDiv scores with average enrichment scores for repetitive elements marked in H3K9me3 datasets from MEF CUT&Tag, MEF CUT&RUN, and mESC CUT&Tag. Number of repetitive elements throughout the entire genome are depicted with various sized points. (C) Profile plots of H3K9me3 MEF CUT&RUN and H3K9me3 mESC CUT&Tag datasets over all RLTR6B, RLTR6-int, IAPEz-int, and L1Md_F2 elements. (D) Profile plots of H2A.Z, H3K27ac, and H3K27me3 CUT&Tag datasets over all RLTR6B, RLTR6-int, IAPEz-int, and L1Md_F2 elements. (E) Genome browser enrichment profile of H2A.Z MEF CUT&Tag, H2A.Z MEF ChIP-Seq, H3K9me3 MEF CUT&Tag, H3K9me3 MEF ChIP-Seq, H3K9me3 MEF CUT&RUN, and H3K9me3 mESC CUT&Tag.

### Many Factors Traditionally Thought to Bind Euchromatin Co-Purify with Insoluble Heterochromatin

Having demonstrated a clear under-representation of heterochromatic repetitive elements within ChIP-based assays (**Fig 6**), we next speculated that proteins bound at heterochromatin loci might be unknowingly excluded from downstream analyses. To investigate this possibility, we prepared crosslinked and sonicated chromatin in a manner similar to the aforementioned mock ChIP-Seq experiments, but rather than investigating the DNA portion of supernatant and pellet fractions, we performed mass spectrometry and identified enriched proteins. Here, we identified 834 soluble proteins significantly enriched in the supernatant (p-value < 0.05, Log2FC > 0.5) and 1509 protein significantly enriched in the insoluble pellet (**Fig 7A**). Intriguingly, gene ontology (GO) analysis revealed an enrichment for proteins involved in nucleic acid binding and chromatin modification in the pellet-enriched fraction, while transmembrane and transporter-associated proteins tended to be enriched in the supernatant (**Fig 7B** and **Supp Tables 1 & 2**). Further inspection revealed several proteins with known function in the centromere or nucleolus to be enriched within the pellet fraction (**Fig 7C & 7D**), likely due to the highly compact nature of these separate nuclear compartments/structures^35,36^. Several zinc-finger family proteins, which are known to function in heterochromatin binding and repetitive element silencing, were also enriched within pellet samples (**Fig 7E**)^37,38^. In addition to these somewhat expected results, we identified many enriched factors involved in epigenetic silencing or transcriptional activation within the pellet fraction. These included well established silencing factors, such as ATRX, DNMT1, DNMT3A, SIRT6, and UHRF1^39,40^, as well as several factors typically thought to function in gene activation and reside within euchromatin, such as BRD4, JMJD6, KAT2B, and NSD1/2 (**Fig 7F**)^41–43^. Perhaps most surprisingly, many well-studied transcription factors with known binding capacity at gene regulatory regions were found to be significantly enriched in the pellet (**Fig 7G**), including ELF1, YY1, RUNX4, and ETV6^44–47^. Taken together, these results indicate that several commonly studied proteins, including several epigenetic components and transcription factors that are traditionally studied in the context of genic euchromatin, are depleted from ChIP-based assays and may have unknown auxiliary functions within heterochromatic portions of the genome.

**Figure 7.**
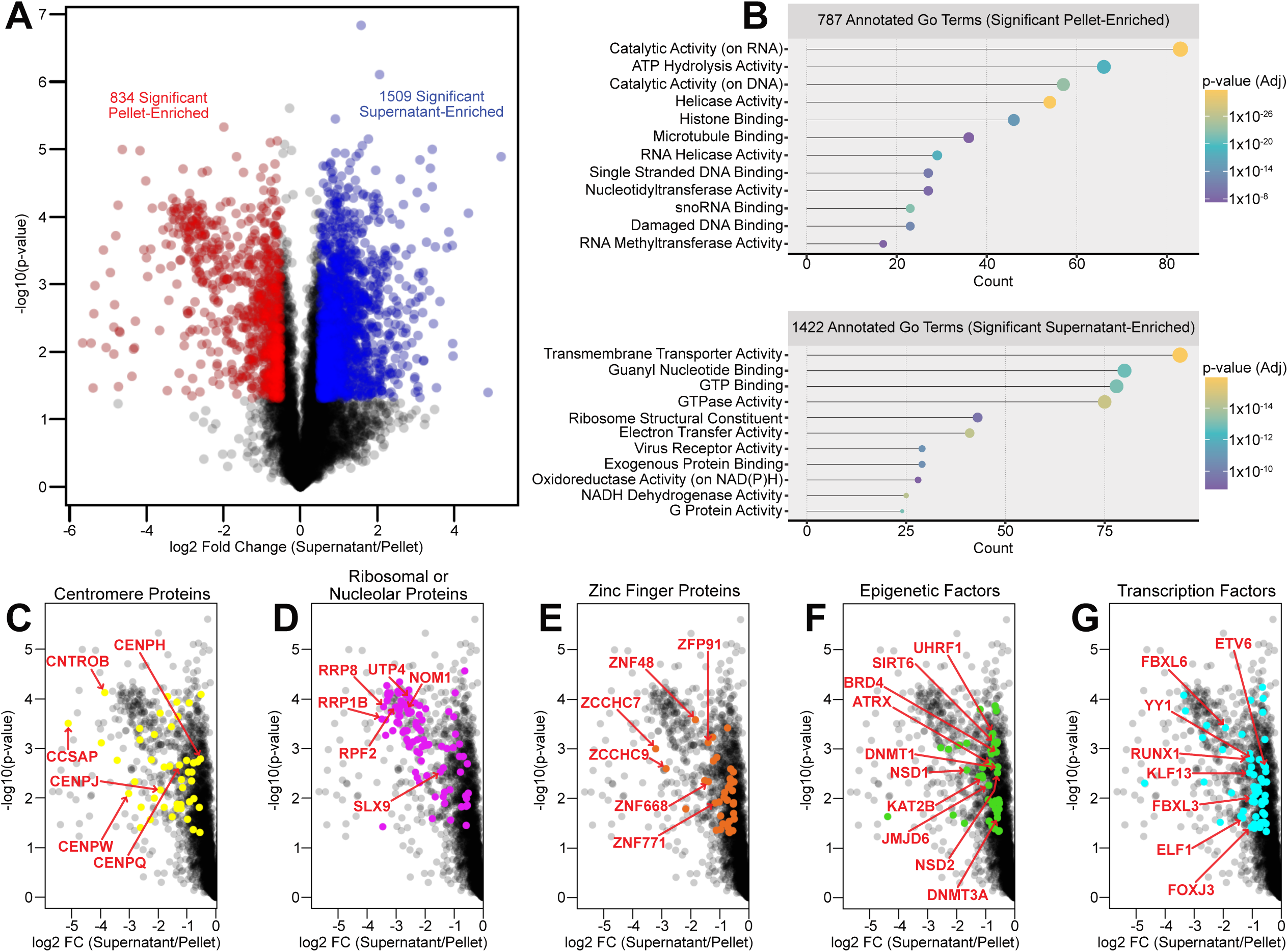
Pelleted Fraction from Crosslinked and Sonicated Samples Contains Well Known Euchromatic Factors. (A) Scatter plot showing significant pellet-enriched (red) and supernatant-enriched (blue) proteins from a mass spectrometry experiment. (B) Gene ontology analysis of the annotated and significant pellet-enriched and supernatant-enriched proteins. (C) Scatter plot showing centromere associated proteins (yellow) that were significantly enriched in the pellet fraction. (D) Scatter plot showing ribosomal and nucleolus associated proteins (purple) that were significantly enriched in the pellet fraction. (E) Scatter plot showing zinc finger proteins (orange) that were significantly enriched in the pellet fraction. (F) Scatter plot showing epigenetic factors (green) that were significantly enriched in the pellet fraction. (G) Scatter plot showing transcription factors (cyan) that were significantly enriched in the pellet fraction.

## Discussion

As proposed in prior studies^10,11^, we find ChIP-based strategies to be biased towards accessible regions of the genome. Since we did not observe such biases in datasets generated by CUT&Tag and CUT&RUN, which utilize *in situ* enzymatic methods to fragment chromatin, it is plausible that the shortcomings of ChIP are due to chromatin purification, crosslinking, and sonication steps^6,31^. It is noteworthy that biases of ChIP-Seq seem to be marginal (and/or mitigated by input normalization) when interrogating activating chromatin modifications, such as H3K27ac, which exhibited similar enrichment patterns for both CUT&Tag and ChIP-Seq datasets in our analyses. Unlike open and accessible genomic regions, the vast majority of loci enriched for H3K9me3 exhibited highly dissimilar enrichment patterns when we compared data generated from ChIP-Seq with CUT&Tag. Since we found that chromatin within the discarded pellet of ChIP samples tended to have higher levels of H3K9me3, as measured by CUT&Tag, we find it likely that many repetitive elements and retrotransposons are missed in many published ChIP-Seq studies, potentially because repetitive loci are more compacted, and thus less sensitive to sonication. These inferences align with previous reports that genomic regions containing H3K9me3 are somewhat resistant to sonication^11^.

While most repetitive elements in the genome are bound by silencing factors, preventing their expression and subsequent spread throughout the genome, at particular times during development a subset of elements, including evolutionarily young retrotransposons, can function as transcriptional regulatory elements and potentially influence proximal gene expression patterns ^16,19^. Here, we demonstrate that CUT&Tag overcomes biases of ChIP-Seq strategies and allows for the investigation of chromatin modifications at what would otherwise be undetectable repetitive regions. These results indicate that our current understanding of chromatin regulation at repetitive elements, or even repetitive element function, may be severely limited. Our measurements of ChIP enrichment discrepancies focused mainly on the repressive mark H3K9me3, which is typically present at silent repetitive elements, but we also observed the presence of H2A.Z at IAPEz-int elements. Whether additional chromatin features that are typically associated with euchromatin (such as H2A.Z) are also bound at repetitive loci remains an intriguing and unexplored possibility. Prior studies have indicated that chromatin modifications such as H3K27ac and H3K4me3 can function in the activation of certain repetitive loci ^16,19^, but it remains unknown how widespread or common this type of regulation takes place. Subsequent research studies are necessary to address this unknown.

As a scientific community, our current understanding of repetitive element regulation and function, as well as protein binding with heterochromatin, has been largely gleaned from decades of ChIP-based studies. With further adoption of *in situ* chromatin fragmentation methods, we now have the opportunity to expand the knowledge base from which new hypotheses, mechanisms, and models are formulated. We find our mass spectrometry results to be particularly interesting in this regard. While we did identify several proteins with known heterochromatic function within the pellet fraction of our experiment, such as DNMT1 and SIRT6, we also uncovered numerous factors that are not known to bind heterochromatin or influence its transcription, including KAT2B, BRD4, and RUNX4. It is quite possible that many of the proteins we identified within the pellet fraction are depleted from ChIP studies, especially when bound to insoluble portions of the genome. Thus, the function of these seemingly euchromatic factors within heterochromatin has remained unknown – due to technical limitations. It is our hope that future researchers take note of the ChIP biases we uncovered and revisit the function of proteins that were previously considered to function exclusively within euchromatin loci.

For the vast majority of prior studies which investigated genomic patterns of chromatin features, ChIP-Seq has been the preferred method. Consequently, our results suggest that much of what we know about chromatin regulation over repetitive elements is incomplete, and many unknown factors could be involved in repetitive element or heterochromatin regulation. In addition to extending our knowledge of basic mechanisms, further investigation of repetitive loci using *in situ* methods could have translational impacts, in the context of both development and disease. For example, abnormal H3K9me3 levels have been observed in several cancer types, but the inability to adequately map the landscape of healthy and diseased tissues has made it difficult to precisely determine the role of H3K9me3 in disease^4,48^.

Moreover, the use of CUT&Tag and CUT&RUN should enable the research community to achieve a more complete understanding of repetitive element function, and potentially better target chromatin machinery therapeutically. In addition to expanding the assayable portion of the genome, our study offers an approach that could allow forthcoming researchers to investigate the role of what would otherwise be considered euchromatic proteins within more compacted gene-poor genomic loci. With emerging technologies like CUT&Tag, along with recent efforts to assemble more complete genomes^14,15^, we foresee an impending “golden age” of repetitive element research, which will undoubtedly reveal novel roles for proteins and repetitive elements in a wide range of critical biological processes.

## Methods

### Cell Culture

Primary MEFs for all CUT&Tag and CUT&RUN experiments were obtained from embryonic day 13.5 mouse embryos and grown in DMEM supplemented with 10% FBS and 1% penicillin-streptomycin. Cells were cultured at 37°C.

### Antibodies

The following antibodies were used for CUT&Tag experiments: Active Motif #39113 (H2A.Z), Active Motif #39133 (H3K27ac), Active Motif #39155 (H3K27me3), and Active Motif #39161 (H3K9me3). Invitrogen #A6455 was used to target GFP in the Cas-CUT&Tag experiments. Active Motif #39161 was used to target H3K9me3 in the CUT&RUN experiments. Novus Biologicals #NBP 1-72763 was used as the anti-rabbit secondary antibody in all experiments.

### pA-Tn5 Purification and Adaptor Loading

pA-Tn5 was purified and loaded with sequencing adaptors as previously described^6^.

### CUT&Tag

Aliquots of cells were centrifuged at 600xg for 3 minutes at room temperature. Supernatant was decanted, and cellular pellets were resuspended in 400 µL of Nuclear Extraction Buffer (20 mM HEPES-KOH pH 7.9; 10 mM KCl; 0.5 mM spermidine; 0.1% Triton X-100; 20% glycerol; 1x Protease Inhibitor (Pierce #A32963); in autoclaved dH_2_O). Samples were left on ice for 10 minutes and then centrifuged at 1300xg for 4 minutes at 4°C. Supernatant was decanted, and cellular pellets were resuspended in 400 µL of PBS. Samples were centrifuged at 1300xg for 4 minutes at 4°C. Supernatant was decanted, and cellular pellets were resuspended in 1 mL of Wash Buffer (20 mM HEPES-KOH pH 7.5; 150 mM NaCl; 0.5 mM spermidine; 1x Protease Inhibitor (Pierce #A32963); in autoclaved dH_2_O) + 10% DMSO. Samples were placed in a Cryo 1°C Freezing Container (Nalgene #5100-0001) and stored at -80°C until use. Samples were removed from the -80°C freezer and allowed to thaw to room temperature. BioMag Plus Concanavalin A coated magnetic beads (Bangs Laboratories #BP531) were prepared by mixing 10 µL of beads (per sample) with 100 µL of Bead Binding Buffer (20 mM HEPES-KOH pH 7.9; 10 mM KCl; 1 mM CaCl_2_; 1 mM MnCl_2_; in autoclaved dH_2_O). Beads were then placed on a magnetic rack, and supernatant was removed and discarded. Beads were then resuspended in another 1.5 mL of Binding Buffer, then placed on a magnetic rack, and supernatant was removed and discarded. Beads were then resuspended in 10 µL (per sample) of Binding Buffer and held at room temperature until ready to mix with thawed samples. 10 µL of activated beads were added per CUT&Tag sample and incubated at room temperature for 10 min on an end-over-end rotator. Samples were placed on a magnetic rack and supernatant was removed and discarded. Samples were resuspended in 50 µL of Antibody Binding Buffer (Wash Buffer + 0.05% digitonin; 2 mM EDTA; 0.1% BSA) with 1 µL of primary antibody (H2A.Z = Active Motif Cat# 39113; H3K27ac = Active Motif Cat# 39133; H3K27me3 = Active Motif Cat# 39155; H3K9me3 = Active Motif Cat# 39161). Samples were incubated on a nutator overnight at 4°C. Samples were placed on a magnetic rack and supernatant was removed and discarded. Samples were resuspended in 100 µL of Dig-Wash Buffer (Wash Buffer + 0.05% Digitonin) with 1 µL of secondary antibody (Novus Biologicals Cat# NBP1-72763) and incubated on a nutator for 1 hour at room temperature. Samples were placed on a magnetic rack and supernatant was removed and discarded.

While still on the magnetic rack, samples were washed 3 times with 800 µL Dig-Wash Buffer. After 3 washes, supernatant was removed, and samples were resuspended in 100 µL of Dig300 Buffer with 1 µL of pA-Tn5 (157 µg/mL) loaded with sequencing adaptors. Samples were incubated on a nutator for 1 hour at room temperature. Samples were placed on a magnetic rack and supernatant was removed and discarded. While still on the magnetic rack, samples were washed 3 times with 800 µL of Dig300 Buffer (Wash Buffer + 150 mM NaCl; 0.01% Digitonin). After 3 washes, supernatant was removed, and samples were resuspended in 300 µL of Tagmentation Buffer (Dig300 Buffer + 10 mM MgCl_2_). Samples were incubated for 1 hour at 37°C. 10 µL of 0.5M EDTA + 2.5 µL Proteinase K (>600 U/mL, ∼20 mg/mL, Thermo Scientific #EO0491) + 3 µL of 10% SDS were directly added to each sample and mixed by full speed vortexing for 2 seconds. Samples were incubated for 1 hour at 50°C. 300 µL of phenol-chloroform was added to each sample and mixed by full speed vortexing for 2 seconds. Samples were centrifuged at 16,000xg for 3 minutes at room temperature. 300 µL of chloroform was added to each sample and mixed by inverting 10 times. Samples were centrifuged at 16,000xg for 3 minutes at room temperature. The top aqueous layer of each sample was transferred to new Eppendorf tubes containing 750 µL of 100% ethanol + 1 µL of GlycoBlue Coprecipitant (Invitrogen #AM9515, 15 mg/mL) and pipetted up and down to mix. Each sample was chilled on ice for 3 minutes before centrifuging at 16,000xg for 15 minutes at 4°C. Supernatant was decanted and the remaining pellet was washed in 1 mL of 100% ethanol. Samples were centrifuged at 16,000xg for 1 minute at 4°C. Supernatant was removed with a pipette and samples were allowed to air dry completely (approximately 5 minutes). Each pellet was resuspended in 25 µL of RNase Solution (400 µL of autoclaved dH_2_O + 1 µL of RNase A (20 mg/mL, PureLink #12091-021)) and incubated for 10 minutes at 37°C. Purified DNA samples were then stored at -20°C until PCR amplification and sequencing.

### CUT&RUN

CUT&RUN experiments were conducted using the Epicypher protocol, as previously described at (https://www.epicypher.com/content/documents/protocols/cutana-cut&run-protocol-2.1.pdf).

### Preparing Sonicated MEFs for Sequencing

MEFs were grown to confluency in a 10 cm plate. Media was removed and cells were washed with 5 mL of PBS. To crosslink, cells were treated with 1% paraformaldehyde (Pierce #28906) in PBS for 10 minutes at room temperature. To stop crosslinking, 125 mM glycine was added to each plate. Cells were harvested with a cell scraper and washed with 5 mL of PBS. Cells were suspended in 2 mL of 1% SDS Lysis Buffer (83 mM Tris-HCl; 167 mM NaCl; 1.1% Triton X-100; 0.05% SDS; 1x Protease Inhibitor (Pierce #A32963); in autoclaved dH_2_O) and allowed to incubate at room temperature for 10 minutes. Cells from each confluent plate were then equally divided into 4 Eppendorf tubes. Samples were then sonicated for 0, 1, 2, or 3 cycles (pulse = 10s; rest = 20s; amplitude = 30%; 5 min on), keeping tubes on ice between cycles. Samples were centrifuged at 3,000xg for 10 minutes at room temperature and then separated into supernatant and pellet fractions. 5 µL of Proteinase K (>600 U/mL, ∼20 mg/mL, Thermo Scientific #EO0491) + 20 mM EDTA was added to each supernatant sample, and each pellet was resuspended in 500 µL of 1% SDS Lysis Buffer + 5 µL of Proteinase K (>600 U/mL, ∼20 mg/mL, Thermo Scientific #EO0491) + 20 mM EDTA. Pellet samples were then broken up with a 20-gauge syringe. All supernatant and pellet samples were vortexed to mix and incubated for 1 hour at 50°C. 1% SDS was added to each sample, and the pellet samples were again broken up with a 20-gauge syringe. All samples were incubated overnight at 65°C. 300 µL of phenol-chloroform was added to each sample and mixed by full speed vortexing for 30 seconds. Samples were centrifuged at 16,000xg for 3 minutes at room temperature. 300 µL of chloroform was added to each sample and mixed by full speed vortexing for 30 seconds. Samples were centrifuged at 16,000xg for 3 minutes at room temperature. The top aqueous layer of each sample was transferred to new Eppendorf tubes containing 750 µL of ice cold 100% isopropanol + 1 µL of GlycoBlue Coprecipitant (Invitrogen #AM9515) and pipetted up and down to mix. Each sample was chilled on ice for 3 minutes before centrifuging at 16,000xg for 15 minutes at 4°C. Supernatant was decanted and the remaining pellet was washed in 1 mL of 100% ice cold ethanol. Samples were centrifuged at 16,000xg for 5 minutes at 4°C. Supernatant was removed with a pipette and samples were allowed to air dry completely (approximately 5 minutes). Each pellet was resuspended in 25 µL of RNase Solution (400 µL of autoclaved dH_2_O + 1 µL of RNase A (20 mg/mL, PureLink #12091-021)) and incubated for 30 minutes at 37°C. Tubes were briefly flicked to mix the samples and then incubated for another for 30 minutes in a heat block set to 37°C. Purified DNA samples were then stored at -20°C until sequencing adaptors were added.

### Adding Sequencing Adaptors to X-Linked Supernatant Samples

In a PCR strip tube, 10 ng of purified supernatant DNA was mixed with dH_2_O up to 25 µL. 3.5 µL of NEBNext Ultra II End Prep Reaction Buffer and 1.5 µL of NEBNext Ultra II End Prep Enzyme Mix were added to each tube and pipetted to mix (NEBNext Multiplex Oligos for Illumina #E7600S). Samples were placed in a thermocycler to amplify DNA (Lid = 60°C; 20°C for 30 minutes; 65°C for 30 minutes; hold at 4°C). 15 µL of NEBNext Ultra II Ligation Master Mix and 0.5 µL of NEBNext Ligation Enhancer were added to each sample. 1.25 µL of NEBNext i5 and i7 Adaptors (diluted 1:10 in dH2O) were added to each sample and immediately pipetted to mix. Tubes were incubated in a thermocycler for 15 minutes at 20°C (heated lid off). 1.5 µL of USER enzyme was added to each sample (NEB #E7602A). Samples were mixed well and incubated in a thermocycler for 15 minutes at 37°C (lid = 47°C). SPRIselect beads (Beckman Coulter Inc #B23317) were used to clean up samples using the manufacturer’s protocol (1.0x volume) and the final volume (∼13 µL) was transferred to a new clean tube. 15 µL of NEBNext Ultra II Q5 Master Mix was added to each tube, and DNA was amplified using i5 and i7 PCR primers in a thermocycler (98°C for 45 seconds; 14 cycles of 98°C for 15 seconds + 60°C for 10 seconds; 72°C for 1 minute). DNA samples were again cleaned up with SPRIselect beads (Beckman Coulter Inc #B23317) using the manufacturer’s protocol (1.0x volume) and the final volume (∼13 µL) was transferred to a new clean tube. Samples were stored at -20°C until sequencing.

### Adding Sequencing Adaptors to X-Linked Pellet Samples

All purified pellet DNA was combined with 25 µL of 2X Tagmentation Buffer (20 mM Tris; 10 mM MgCl_2_; 5% dimethylformamide; 66% PBS; 0.2% Tween20; in autoclaved dH_2_O) + 1 µL Tn5 + autoclaved dH_2_O up to 50 µL. Samples were incubated at 37°C for 30 minutes at 1000 RPM. 0.2% SDS was added to each tube and samples were incubated at room temperature for 5 minutes. Samples were cleaned up with SPRIselect beads (Beckman Coulter Inc #B23317) using the manufacturer’s protocol (1.1x volume) and the final volume (∼24 µL) was transferred to a new clean tube. 21 µL of purified pellet DNA was mixed with 25 µL of NEBNext High-Fidelity 2X PCR Mastermix (NEB #M0541S), and DNA was amplified using i5 and i7 PCR primers in a thermocycler (72°C for 5 minutes; 98°C for 30 seconds; 13 cycles of 98°C for 10 seconds + 63°C for 15 seconds; 72°C for 1 minute; hold at 4°C). Samples were cleaned up with SPRIselect beads using the manufacturer’s protocol (1.1x volume) and the final volume (∼24 µL) was transferred to a new clean tube. Samples were stored at -20°C until sequencing.

### Library Preparation and Sequencing Data

To amplify the CUT&Tag libraries from various cell lines and ChIP input libraries from sonicated MEFs, 21 µL of purified DNA was mixed with 25 µL NEBNext HiFi 2× PCR Master mix, and 2 µL of unique i5 and i7 barcoded primers, giving a different barcode to each sample. CUT&Tag and ChIP input samples were pooled and sequenced either by NovoGene or the UR-Genomics Research Center, using short-read Illumina next generation sequencing platforms. Raw and processed sequencing data generated in this study can be found at NCBI GEO with the accession number (GSE…).

### Mass Spectrometry

MEFs were grown to confluency in a 10 cm plate. Media was removed and cells were washed with 5 mL of PBS. To crosslink, cells were treated with 1% paraformaldehyde (Pierce #28906) in PBS for 10 minutes at room temperature. To stop crosslinking, 125 mM glycine was added to each plate. Cells were harvested with a cell scraper and washed with 5 mL of PBS. Cells were suspended in 2 mL of 1% SDS Lysis Buffer (83 mM Tris-HCl; 167 mM NaCl; 1.1% Triton X-100; 0.05% SDS; 1x Protease Inhibitor (Pierce #A32963); in autoclaved dH_2_O) and allowed to incubate at room temperature for 10 minutes. Cells from each confluent plate were then equally divided into 4 Eppendorf tubes. Samples were then sonicated for 2 cycles (pulse = 10s; rest = 20s; amplitude = 30%; 5 min on), keeping tubes on ice between cycles.

Samples were centrifuged at 3,000xg for 10 minutes at room temperature and then separated into supernatant and pellet fractions. 20 mM EDTA was added to each supernatant sample, and each pellet was resuspended in 500 µL of 1% SDS Lysis Buffer + 20 mM EDTA. Pellet samples were then broken up with a 20-gauge syringe. All supernatant and pellet samples were vortexed to mix and incubated for 1 hour at 50°C. 1% SDS was added to each sample, and the pellet samples were again broken up with a 20-gauge syringe. 200 µM NaCl was added to all samples to reverse crosslinks, and all samples were incubated overnight at 65°C. Pellet samples were again broken up with a 20-gauge syringe.

Samples were concentrated by adding 6x volumes of ice-cold acetone and incubating for 30 minutes. Samples were centrifuged at 15,000xg for 5 minutes. Supernatant was discarded and samples were air dried for 5 minutes. Samples were then solubilized and run on a 4-12% SDS-PAGE gel. The gel was stained with SimplyBlue SafeStain (Invitrogen) and washed overnight. Gel slices were excised, cut into 1mm cubes, and destained. The destained gel slices were reduced with DTT (Sigma) and alkylated with IAA (Sigma), and then dehydrated with acetonitrile. Trypsin (Promega) was diluted to 10 ng/µL in 50 mM ammonium bicarbonate and used to cover the dehydrated gel slices. The slices were incubated in the trypsin for 30 minutes at room temperature. Additional ammonium bicarbonate was added until the gel pieces were completely submerged, and the gel pieces were then incubated overnight at 37°C. The next day, peptides were extracted from the gel slices by adding 50% acetonitrile and 0.1% TFA, and then dried using a CentriVap concentrator (Labconco). Desalting was performed with homemade C18 spin columns, followed by drying, and reconstitution in 0.1% TFA. A fluorometric peptide assay (Thermo Fisher) was used to determine the final peptide concentrations.

The extracted peptides were then used for mass spectrometry experiments. Peptides were injected onto a 75 µm x 2 cm trap column (Thermo Fisher) and then refocused on an Aurora Elite 75 µm x 15 cm C18 column (IonOpticks) using a Vanquish Neo UHPLC (Thermo Fisher) attached to an Orbitrap Astral mass spectrometer (Thermo Fisher). Solvent A used for these experiments was 0.1% formic acid in water, and solvent B was 0.1% formic acid in 80% acetonitrile. Ions were added to the mass spectrometer with an Easy-Spray source operating at 2 kV. The solvent gradient started at 1% solvent B and increased to 5% solvent B over 0.1 minutes. The solvent gradient further increased to 30% solvent B in 12.1 minutes, 40% solvent B in 0.7 minutes, and finally 99% solvent B in 0.1 minutes. The gradient was held at 99% solvent B for 2 minutes to wash the column (total runtime 15 minutes). The column was re-equilibrated with 1% solvent B between each mass spectrometry run. The Orbitrap Astral was used in data-independent acquisition (DIA) mode, and MS1 scans were acquired in the Orbitrap at a resolution of 240,000. The maximum injection time was 5 ms covering a range of 380-980 m/z. DIA MS2 scans were acquired in the Astral mass analyzer using a 6 ms maximum injection time with variable windowing (4 Da from 380-750 m/z and 6 Da from 750-980 m/z). The HCD collision energy was 28%, and the normalized AGC was 500%. Fragment ions were acquired from 150-2000 m/z with a cycle time of 0.6 seconds.

### Bioinformatic Analysis

Raw mass spectrometry data were processed using DIA-NN version 1.8.1 (https://github.com/vdemichev/DIA-NN) using library-free analysis mode^49^. The *Mus musculus* UniProt ‘one protein sequence per gene’ database (UP000000589_10009, downloaded 4/7/2021) was used to annotate the dataset while enabling ‘deep learning-based spectra and RT prediction’. Precursor ion generation settings included a maximum of 1 missed cleavages, a maximum of 1 variable modifications for Ox(M), a peptide length range of 7-30, a precursor charge range of 2-4, a precursor m/z range of 380-980, and a fragment m/z range of 150-2000. Quantification was performed with ‘Robust LC (high precision)’ mode, using RT-dependent normalization, MBR enabled, protein inferences set to ‘Genes’, and ‘Heuristic protein inference’ turned off. Mass tolerances and scan window sizes were automatically determined by the software. Precursors were filtered at a library precursor q-value of 1%, a library protein group q-value of 1%, and a posterior error probability of 50%. Protein quantification was performed using the MaxLFQ algorithm in the DIA-NN R package (https://github.com/vdemichev/diann-rpackage). The number of peptides in each protein group was counted with the DiannReportGenerator Package (https://github.com/URMC-MSRL/DiannReportGenerator)^50^.

Publicly available datasets were downloaded from ENA. CutAdapt was used to trim the adaptor sequences from CUT&Tag datasets with parameters -m 1 -a CTGTCTCTTATA -A CTGTCTCTTATA. Fasta files were aligned to the mouse (mm10) and human (hg38) genomes with Bowtie2. PICARD MergeSamFiles was used to convert .sam files to .bam files with parameters SO= coordinate CREATE_INDEX=true. PICARD MarkDuplicates was used to remove duplicate reads from all .bam files with parameters REMOVE_DUPLICATES=true CREATE_INDEX=true. Deeptools BAMcoverage was used to convert .bam files to .bw files with parameters --normalizeUsing RPKM --binSize 10 –extendReads 100. UCSC bigwigtobedgraph was used to convert .bw files to .bedgraph files. UCSC bigWigMerge was used to merge all replicates from each experiment MACS2 bdgcmp was used to calculate ChIP-Seq enrichment scores above background input levels with parameters -m ppois. MACS2 bdgcmp was also used to calculate enrichment scores of pellet and supernatant samples over one another with parameters -m logFE -p 10. MACS2 bdgpeakcall was used to call peaks on all datasets with parameters -g 100 -l 100. Various -c values were used with MACS2 bdgpeakcall to generate roughly 10k or 30k peaks, and the resulting peak sets were trimmed to exactly the top 10k or 30k locations in R based on RPKM or ppois enrichment scores. For S1-specific and P1-specific peaksets, MACS2 bdgpeakcall was used with parameters -g 100 -l 100 and c = 0.4 on the S1/P1 and P1/S1 bdgcmp files. Genome browser enrichment profiles were generated with IGV. HOMER annotatePeaks was used to determine genomic annotations for the most highly enriched regions in each dataset, as well as distance to nearest TSS and CpG density using parameters -CpG and mm10 or hg38 genomes downloaded from HOMER. Deeptools multiBigwigSummary was used with parameters BED-file and --outRawCounts to calculate enrichment scores that were then assigned a rank, and ranks were used along with the pHeatmap R package to generate rank-normalized heatmaps with the parameters cluster_rows= FALSE, cluster_cols = FALSE, col = colorRampPalette(c(“lightblue”, “darkblue”))(256). Deeptools multiBigwigSummary was also used with parameters BED-file and --outRawCounts to calculate enrichment scores that were used in making scatter plots and RPKM boxplots in R. Scatter plots were made using the R packages ggplot2 and ggpointdensity with the parameters geom_pointdensity(alpha=0.1, size = 3) + scale_color_gradient(low=“#041370”, high=“#FFFF00”) + theme_bw() + theme(panel.grid.major = element_blank(), panel.grid.minor = element_blank(). Deeptools plotHeatmap was used to generate standard heatmaps using parameters --missingDataColor white --colorList “white,red,maroon,purple” -- yMin 0, as well as desired –yMax and –zMax values. Bedtools intersect was used to determine overlapping regions of datasets with parameters -wa | uniq . The dplyr and scales R packages were used to filter datasets by repeat name, as well as calculate number of repeats, average milliDiv score, and average ranks for each repeat family using parameters filter, group_by, and summarise. plot was used in R to generate rank score difference scatter plots with parameters pch=16, col=rgb(0,0,0,0.2), cex=AdjN. Adjusted N values were calculated based on number of repeats calculated by dplyr, with n < 10 = 0.1, 10<n<200 = n/100, and n<200 = 2. LTRs were labeled with points in R using parameters pch=16, col=rgb(0,0,1,0.6), cex=AdjN. LINEs were labeled with points in R using parameters pch=16, col=alpha(“darkorange”, 0.6), cex=AdjN. HOMER analyzeRepeats was used with parameters mm10 - count exons -condenseGenes to calculate scores in Supp Fig 5B. Scores were then given a rank, and normalized to copy # and repeat length before being plotted in R with parameters pch=16, col=rgb(0,0,0,0.2), cex= 0.45. featureCounts in the Rsubread package was used to generate the counts for the average milliDiv vs log10(Average Reads per Copy) plots using the parameters -O -p and -a (RepeatMasker), and then adding a pseudocount of 1. Gene ontology analysis was conducted using Gene IDs for proteins that were significantly (p.value < 0.05) enriched in the pellet or supernatant fractions (log2 fold change > 0.5). Analysis was conducted in R using the clusterProfiler package, with parameters OrgDb = “org.Hs.eg.db”, ont = “MF”, readable = TRUE, fun = enrichGO, qvalueCutoff = 0.05. Gene ontology plots were produced in R with the ggplot2 package with parameters aes(Count, fct_reorder(Description, Count))) + facet_grid(“∼Cluster”) + geom_segment(aes(xend=0,yend=Description)) + geom_point(aes(color=p.adjust, size=GeneRatio*100)) + scale_color_gradientn(colours=c(“#f7ca64”, “#46bac2”, “#7e62a3”), trans=“log10”, guide = guide_colorbar(reverse = TRUE, order = 1)) + theme(panel.border = element_blank(), panel.grid.major = element_line(linetype = ‘dotted’, colour = ‘#808080’), panel.grid.major.y = element_blank(), panel.grid.minor = element_blank(), axis.line.x = element_line()) + scale_size_continuous(range=c(1,5)) + guides(size = guide_legend(override.aes = list(shape=1))).

## Supporting information

SubFigsCombined

SuppTable1

SuppTable2

## Acknowledgements

This work was funded by grants from the National Institute of Health, R35GM137833 (PJM) and R35GM133462 (MRO).

## Author Contributions

CUT&Tag datasets were generated by KC, SH, KM, and MA. CUT&RUN datasets were generated by EC. Mock ChIP-Seq input pellet and supernatant datasets were generated by BP and RL. Data analysis was done by BP. PV provided conceptual guidance throughout the project. The initial manuscript was drafted by BP. Edits to the manuscript were made by PJM and MRO. The entire project was jointly supervised by PJM and MRO.

