## Supplementary material for "CUT&Tag Identifies Repetitive Genomic Loci that are Excluded from ChIP Assays": SubFigsCombined

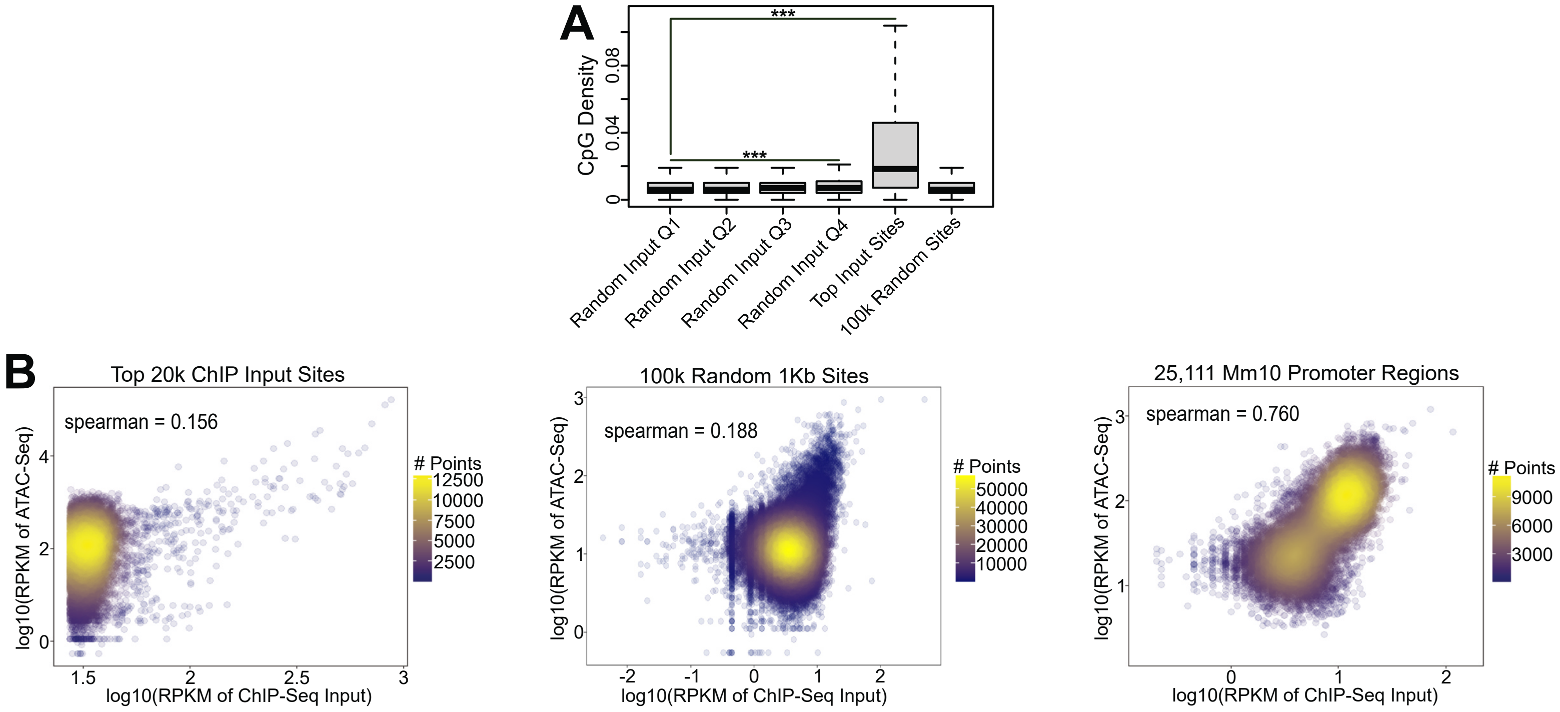


**Figure S1 ChIP-Seq Input is Correlated with ATAC-Seq**

(A) CpG density for ChIP-Seq input (100k random 1Kb regions divided into quartiles based on input enrichment levels), top 20k ChIP-Seq input sites, or 100k random 1Kb sites.

(B) Scatterplots comparing $\log_{10}$(RPKM) scores for ChIP-Seq input and ATAC-Seq at the top 20k ChIP-Seq input sites, 100k random 1Kb sites, and all annotated mouse promoters.

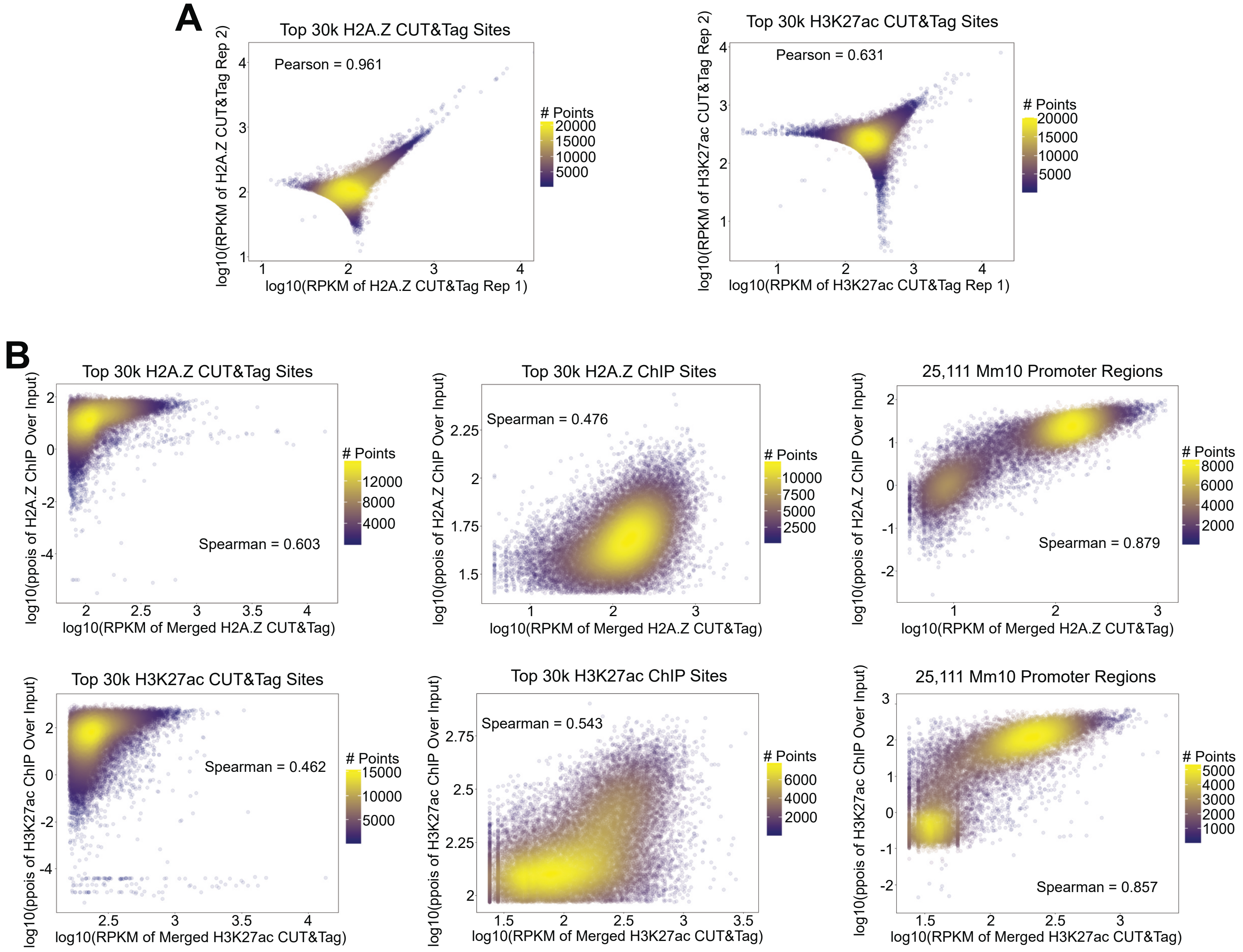


**Figure S2 ChIP-Seq and CUT&Tag Datasets are Correlated for H2A.Z and H3K27ac**

(A) Scatterplots comparing $\log_{10}$(RPKM) scores of individual replicates of H2A.Z and H3K27ac CUT&Tag datasets over the most highly enriched regions.

(B) Scatterplots comparing enrichment scores of H2A.Z CUT&Tag with H2A.Z ChIP over the top 30k H2A.Z CUT&Tag sites, the top 30k H2A.Z ChIP-Seq sites, and all annotated mouse promoters, and H3K27ac CUT&Tag with H3K27ac ChIP-Seq over the top 30k H3K27ac CUT&Tag sites, the top 30k H3K27ac ChIP-Seq sites, and all annotated mouse promoters.

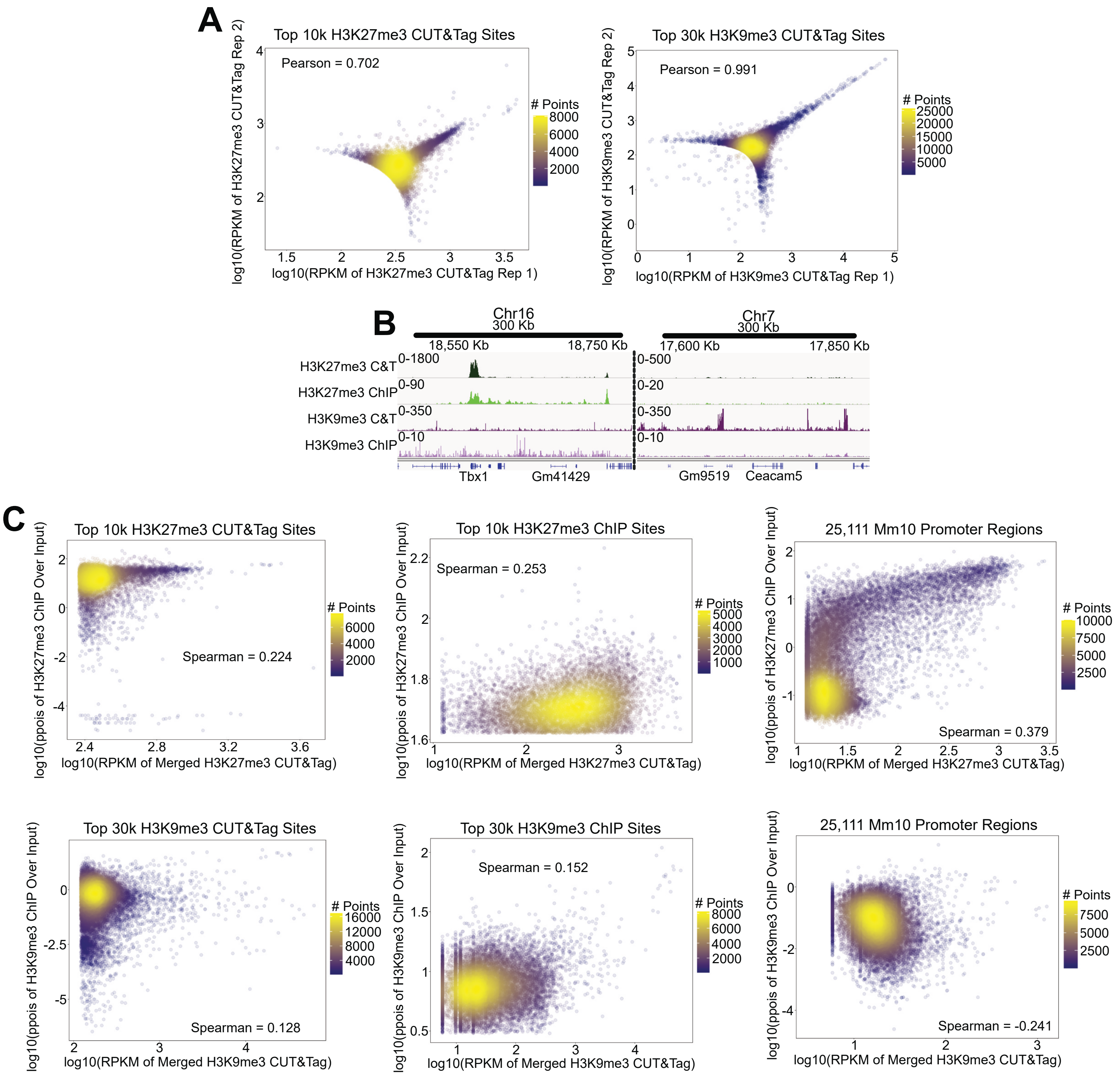


**Figure S3 ChIP-Seq and CUT&Tag Datasets are Correlated for H3K27me3, but Not H3K9me3**

(A) Scatterplots comparing $\log_{10}$(RPKM) scores of individual replicates of H3K27me3 and H3K9me3 CUT&Tag datasets over the most highly enriched regions.

(B) Additional genome browser enrichment profiles of H3K27me3 and H3K9me3 CUT&Tag and ChIP-Seq datasets showing overlap between the methods for H3K27me3, but not H3K9me3.

(C) Scatterplots comparing enrichment scores of H3K27me3 CUT&Tag with H3K27me3 ChIP over the top 10k H3K27me3CUT&Tag sites, the top 10k H3K27me3 ChIP-Seq sites, and all annotated mouse promoters, and H3K9me3 CUT&Tag with H3K9me3 ChIP-Seq over the top 30k H3K9me3 CUT&Tag sites, the top 30k H3K9me3 ChIP-Seq sites, and all annotated mouse promoters.

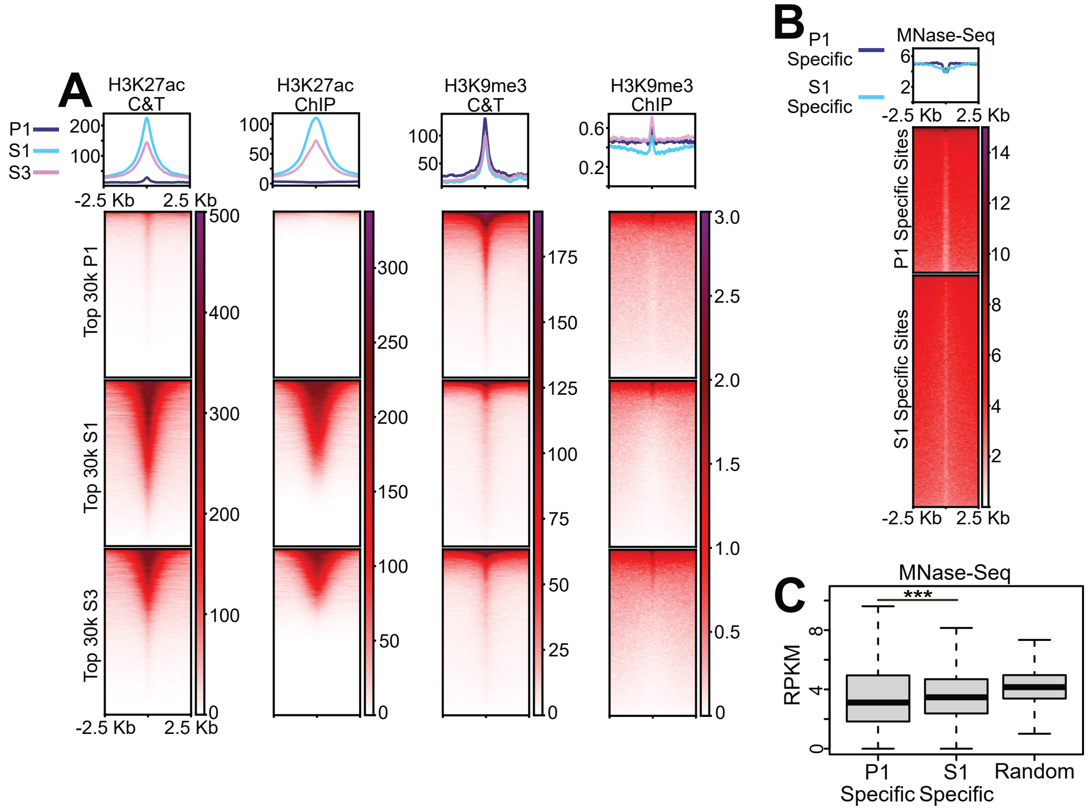


**Figure S4 MNase-Based Genome Fragmentation Techniques Do Not Have the Same Biases as Traditional ChIP**

(A) Heatmaps and profile plots of H3K27ac and H3K9me3 CUT&Tag and ChIP-Seq signal over the 30k most highly enriched P1, S1, and S3 sites.

(B) Heatmap and profile plot of MNase-Seq signal over the most highly enriched P1-specific and S1-specific regions.

(C) Enrichment scores of MNase-Seq signal over the most highly enriched P1-specific and S1-specific regions, and 100k random 1Kb regions.

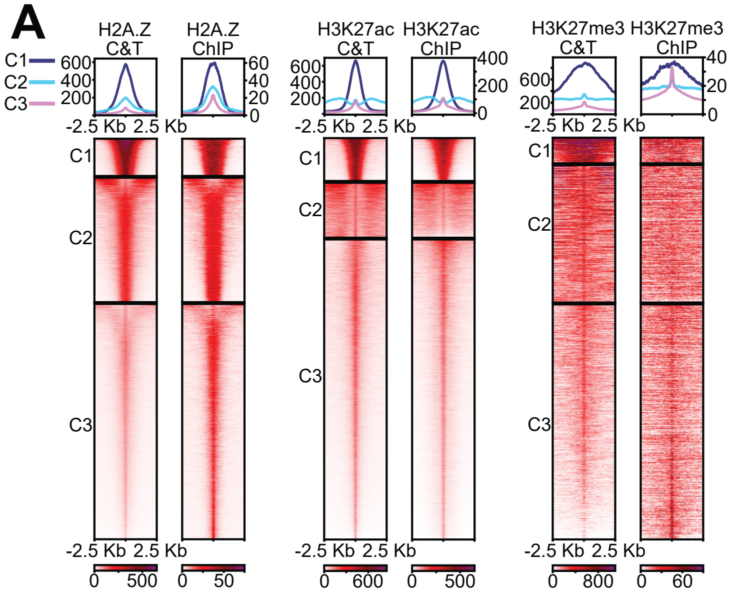


**Figure S5 ChIP-Seq and CUT&Tag Produce Similar Patterns of Enrichment for H2A.Z, H3K27ac, and H3K27me3**

(A) Heatmaps and profile plots of H2A.Z, H3K27ac, and H3K27me3 CUT&Tag and ChIP-Seq datasets over union files of all the most highly enriched CUT&Tag and ChIP-Seq sites, sorted by *k*-means clustering (C1-C3).

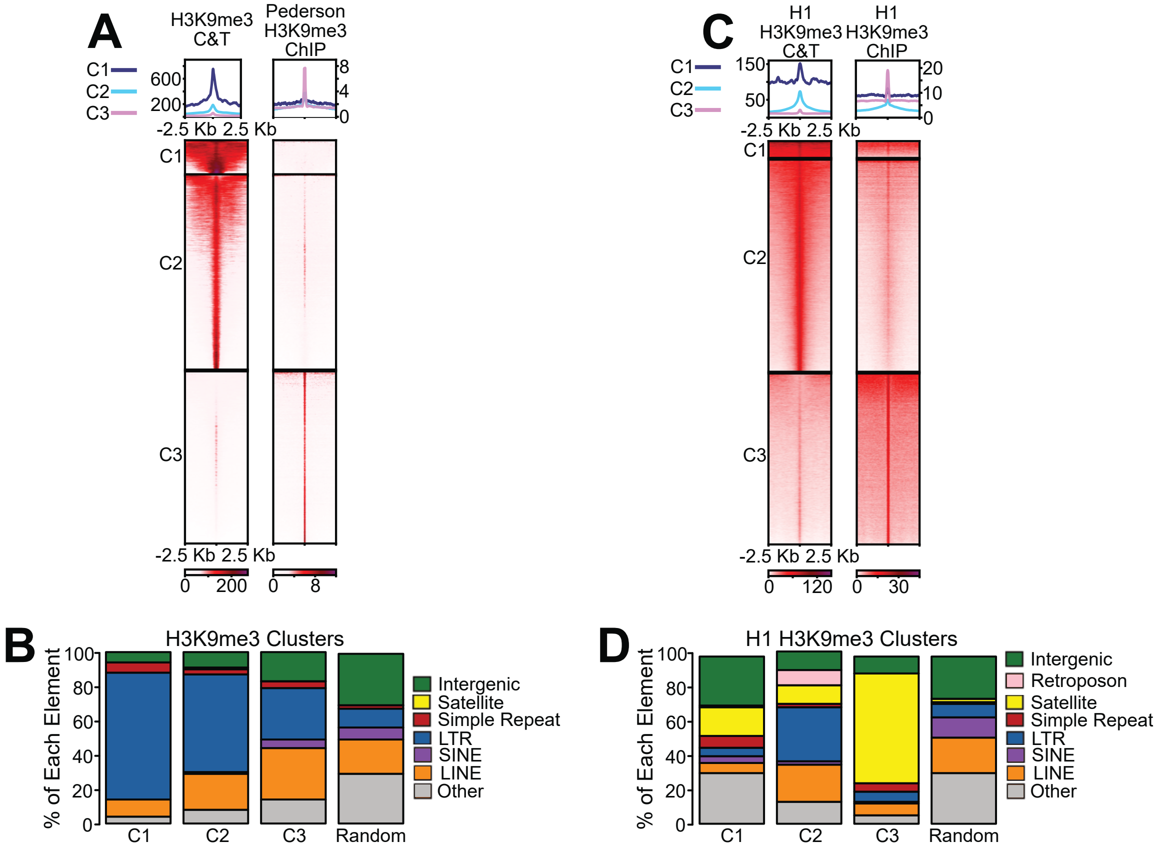


**Figure S6 LTRs are Consistently Enriched in H3K9me3 CUT&Tag Datasets**

(A) Heatmaps and profile plots of MEF H3K9me3 CUT&Tag data and MEF H3K9me3 ChIP-Seq data from Pederson et al. over union files of all the most highly enriched CUT&Tag and ChIP-Seq sites, sorted by *k*-means clustering (C1-C3).

(B) Genomic annotation of the repetitive elements enriched in each MEF H3K9me3 cluster (C1-C3).

(C) Heatmaps and profile plots of H1 H3K9me3 CUT&Tag data and H1 H3K9me3 ChIP-Seq data over union files of all the most highly enriched CUT&Tag and ChIP-Seq sites, sorted by *k*-means clustering (C1-C3).

(D) Genomic annotation of the repetitive elements enriched in each H1 H3K9me3 cluster (C1-C3).
